# A patient-derived blood-brain barrier model for screening copper bis(thiosemicarbazone) complexes as potential therapeutics in Alzheimer’s disease

**DOI:** 10.1101/2023.08.20.554047

**Authors:** Joanna M. Wasielewska, Kathryn Szostak, Lachlan E. McInnes, Hazel Quek, Juliana C. S. Chaves, Jeffrey R. Liddell, Jari Koistinaho, Lotta E. Oikari, Paul S. Donnelly, Anthony R. White

**Affiliations:** Mental Health and Neuroscience Program, QIMR Berghofer Medical Research Institute, Brisbane, QLD, Australia; Faculty of Medicine, University of Queensland, St. Lucia, QLD, Australia; School of Chemistry, Bio21 Institute for Molecular Science and Biotechnology, The University of Melbourne, Parkville, VIC, Australia; School of Biomedical Science, University of Queensland, St. Lucia, QLD, Australia; School of Biomedical Sciences, Faculty of Health, Queensland University of Technology, QLD, Australia; Department of Anatomy and Physiology, The University of Melbourne, Parkville, VIC, Australia; Drug Research Program, Division of Pharmacology and Pharmacotherapy, University of Helsinki, Helsinki, Finland; Neuroscience Centre, Helsinki Institute of Life Science, University of Helsinki, Helsinki, Finland

**Keywords:** Alzheimer’s disease, blood-brain barrier, copper bis(thiosemicarbazone), metal compound, neuroinflammation, drug screening platform, neurotherapeutics

## Abstract

Alzheimer’s disease (AD) is the most prevalent cause of dementia characterised by progressive cognitive decline. Addressing neuroinflammation represents a promising therapeutic avenue to treat AD, however, the development of effective anti-neuroinflammatory compounds is often hindered by their limited blood-brain barrier (BBB) permeability. Consequently, there is an urgent need for accurate, preclinical AD patient-specific BBB models to facilitate the early identification of immunomodulatory drugs capable of efficiently crossing human AD BBB.

This study presents a unique approach to BBB drug permeability screening as it utilises the familial AD patient-derived induced brain endothelial-like cells (iBEC)-based model, which exhibits increased disease relevance and serves as an improved BBB drug permeability assessment tool when compared to traditionally employed *in vitro* models. To demonstrate its utility as a small molecule drug candidate screening platform, we investigated the effects of Cu^II^(atsm) and a library of novel metal bis(thiosemicarbazone) complexes – a class of compounds exhibiting anti-neuroinflammatory therapeutic potential in neurodegenerative disorders. By evaluating the toxicity, cellular accumulation and permeability of those compounds in the AD patient-derived iBEC, we have identified Cu^II^(dtsm) as an emerging drug candidate with enhanced transport across the AD BBB. Furthermore, we have developed a multiplex approach where AD patient-derived iBEC were combined with immune modulators TNFα and IFNγ to establish an *in vitro* model representing the characteristic neuroinflammatory phenotype at the patient’s BBB. Here we observed that treatment with Cu^II^(dtsm) not only reduced the expression of proinflammatory cytokine genes but also reversed the detrimental effects of TNFα and IFNγ on the integrity and function of the AD iBEC monolayer. This suggests a novel pathway through which copper bis(thiosemicarbazone) complexes may exert neurotherapeutic effects in AD by mitigating BBB neuroinflammation and related BBB integrity impairment.

Together, the presented model provides an effective and easily scalable *in vitro* BBB platform for screening AD drug candidates. Its improved translational potential makes it a valuable tool for advancing the development of metal-based compounds aimed at modulating neuroinflammation in AD.

**Graphical abstract:** 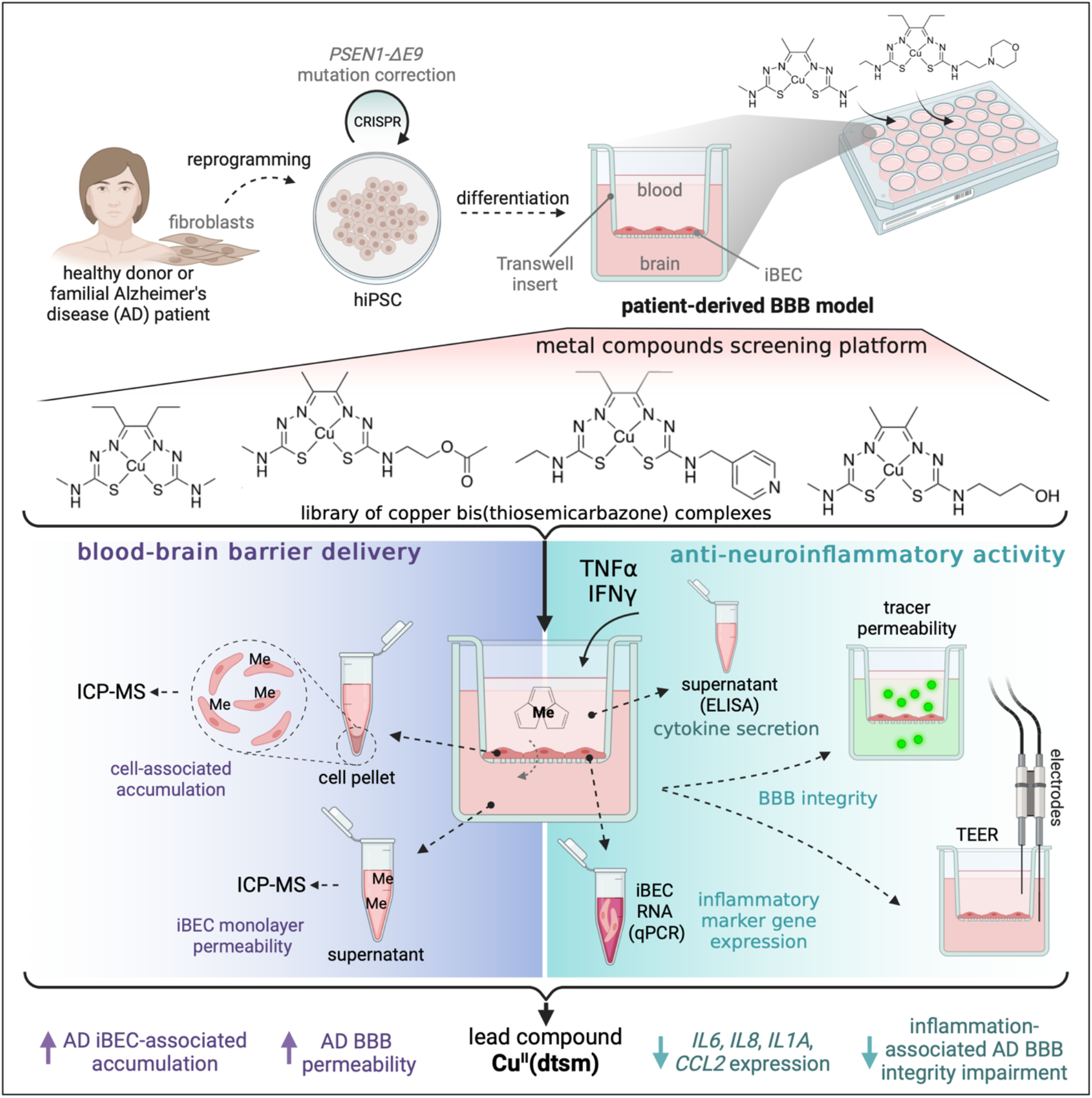

## INTRODUCTION

Alzheimer’s disease (AD) is a progressive neurodegenerative disorder that predominantly manifests as deficits in cognitive functions, such as memory and attention [1]. Recent evidence has shown that neuroinflammation is an important driver of AD, propelling significant research interest towards molecules capable of modulating the immune response within the brain [2], [3]. Yet, despite these efforts, no anti-neuroinflammatory therapeutics have been approved for clinical use in AD.

One of the major hurdles in AD drug development is the blood-brain barrier (BBB) formed by brain endothelial cells (BEC), astrocytes, and pericytes at the blood-brain interface [4]. The BBB is essential for physiological brain function, but restricts the transport of therapeutic agents into the central nervous system (CNS), reducing their clinical effectiveness [5]. Given the importance of improving drug penetration at the level of the BBB, numerous preclinical BBB models have been established and utilised in the AD drug development pipeline [6]–[8]. Although proven useful in some aspects of preclinical drug assessment, those traditionally employed BBB models offer low translational applicability to AD patients, contributing to the modest success rates witnessed in AD clinical trials [9].

An important caveat of conventional cell-monolayer *in vitro* models used in drug permeability screening is their reliance on cell sources that lack clinical relevance. These models frequently utilise human immortalised BEC lines, such as hCMEC/D3, or cells originating from non-CNS, non-endothelial, or non-human cell sources, exemplified by the Caco-2 and MDCK models [10]–[12]. Consequently, those cells demonstrate considerable molecular, phenotypical, and functional differences as compared to human *in vivo* BEC, which can impede the translation of results [13]–[15]. Alternative synthetic systems, such as the parallel artificial membrane permeability assay (PAMPA) model, although continuously modified to achieve higher biomimicry to biological barriers, are deficient in active drug transportation systems and lack the cellular composition of the BBB. Thus, they offer relatively low *in vitro* to *in vivo* drug permeability prediction accuracy [16]–[18].

Concurrently, recent studies have reported on multifactorial BBB dysfunction in AD, suggesting the involvement of BBB cells in disease development and progression [19]–[23]. In addition, neuroinflammation has been shown to drive some of the aspects of BBB impairment in AD linking multiple pathways involved in vascular- and neuro-degeneration [24]–[26]. Identified disease-associated changes at the BBB were also shown to contribute to the development of a complex microenvironment in AD brain barriers with important implications for drug delivery [27]–[30]. Together these observations highlight the key limitation of traditionally used BBB models as they lack disease- and patient-specific characteristics, simultaneously urging the development of accurate BBB *in vitro* screening platforms to reliably model the response of AD patients to drug treatment.

Correspondingly, drug permeability assessed in the modern BBB models derived from healthy donor human induced pluripotent stem cells (hiPSC) was recently shown to closely reflect *in vivo* BBB permeability dynamics in the human brain, suggesting potential high translatability of hiPSC-derived preclinical platforms [31], [32]. The hiPSC-derived induced brain endothelial-like cells (iBEC) generated also a cell monolayer of increased, and hence more physiological, integrity as compared to traditionally used MDCK [33] or Caco-2 [31] models and were suggested to achieve better CNS permeability prediction than PAMPA platforms [34]. In addition, hiPSC can be derived from patient cell sources, with the emerging collection of patient-derived iBEC models now becoming available for various neurodegenerative disorders [22], [35]–[39]. Despite a growing number of patient hiPSC-derived BBB models now being developed and characterised, no reports describe their practical utility in novel anti-neuroinflammatory drug candidate screening in AD.

Our previous studies successfully demonstrated *in vitro* modelling of the BBB from familial AD patients using hiPSC-derived iBEC harbouring a *PSEN1* mutation [40], [41]. These cells exhibited physiologically relevant barrier formation and expressed relevant drug transporters such as P-glycoprotein (*ABCB1*), multidrug resistance protein 1 (*ABCC1*) and breast cancer resistance protein (*ABCG2*) [40], [41]. Here, to enhance the practical application of hiPSC-derived BBB models in the drug discovery pipeline for AD, we sought to validate our AD patient-derived BBB model for testing the barrier-permeability and anti-inflammatory properties of novel neuro-pharmaceuticals.

To that end, we designed a library of metal bis(thiosemicarbazone) (btsc) complexes, incorporating copper or nickel (**Figure 1A**). This library of compounds consisted of structural derivatives of diacetylbis(N(4)-methylthiosemicarbazonato) copper(II) (Cu^II^(atsm)), which has broad therapeutic potential in several preclinical models of neurodegeneration [42]–[48]. We initially assessed the cytotoxicity of the novel metal compounds in human vascular endothelial cells and subsequently, we investigated the toxicity, accumulation and permeability of these compounds across the BBB using both control and familial AD patient-derived iBEC models. Considering that anti-neuroinflammatory actions are now recognised as one of the primary mechanisms underlying neurotherapeutic effects of Cu(btsc) complexes [42], [48], [49], we aimed to investigate the immunomodulatory properties of the tested compounds in our model. To facilitate that, we multiplexed AD patient-derived iBEC with neurologically relevant immune modulators, namely tumour necrosis factor α (TNFα) and interferon γ (IFNγ)[2], resulting in the development of neuroinflammatory phenotype at the patient-derived BBB *in vitro*. By adopting this strategy, we were able to assess the dynamics of drug permeability across the BBB and simultaneously pre-screen their potential anti-inflammatory activity within the same AD patient-cell platform, efficiently identifying promising compounds for further assessment.

**Figure 1.**
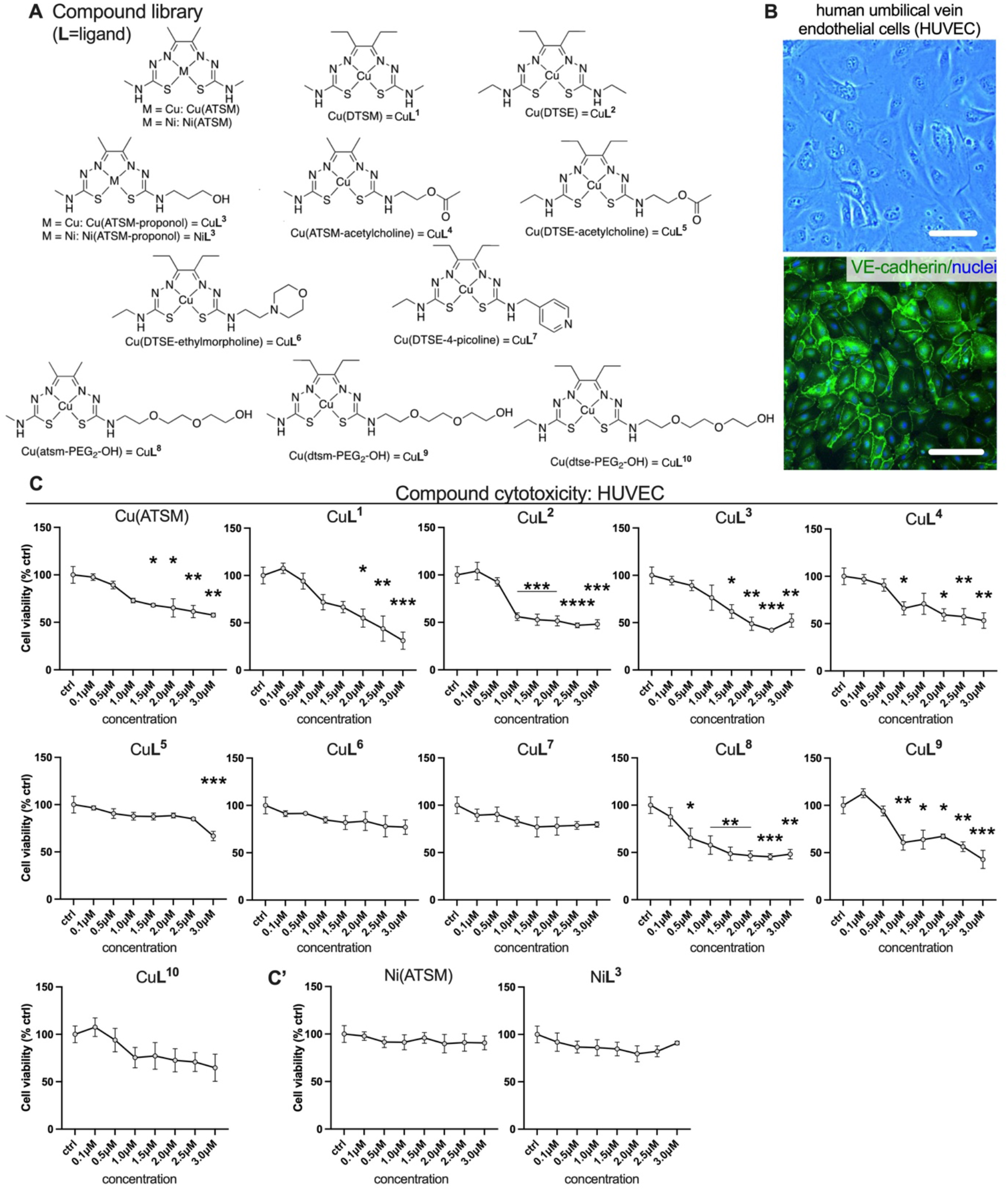
Effects of metal compounds on the viability of the human umbilical vein endothelial cells (HUVEC). (**A**) Chemical structures of Cu(ATSM) and novel metal compounds investigated in the study. (**B**) Representative phase-contrast image (top panel) and immunofluorescence image of vascular endothelial cell marker VE-cadherin (green) with Hoechst nuclear counterstaining (bottom panel) in HUVEC. Scale bar, 100 μm. (**C-C’**) HUVEC viability after treatment with Cu compounds and Ni compounds as assessed with MTT assay. Cell viability is shown as % of viable cells compared to untreated control (ctrl). n=2 for Cu(ATSM) and n=3 independent replicates for other compounds. Data are presented as mean ± SEM. Statistical analysis was performed using one-way ANOVA with Dunnett’s test. \**p*<0.05, \*\**p*<0.01, \*\*\**p*<0.001, \*\*\*\**p*<0.0001. L=ligand.

In summary, this study presents a novel approach to neuro-immunomodulatory drug candidate BBB permeability screening in a familial AD context.

## RESULTS

### Cu^II^ and Ni^II^ form novel complexes with bis(thiosemicarbazone) ligands

To validate the application of our familial AD patient-derived BBB platform as a novel screening tool for small molecule drug candidates, we developed and investigated a library of metal-(btsc) compounds with modifications to its (btsc) ligand (**L**) framework (**Figure 1A**).

Bis(thiosemicarbazone) ligands derived from 1,2-diones undergo double deprotonation and act as dianionic tetradentate N_2_S_2_ ligands to form charge neutral, lipophilic complexes with copper(II). Cu(btsc) complexes are stable with respect to dissociation of the metal (*K_a_* ∼ 10^18^) and are often membrane permeable. The biodistribution, cellular accumulation, and metabolism of Cu(btsc) is dictated by the nature of the substituents on the ligand backbone. These substituents alter lipophilicity, solvation, membrane permeability and the interaction with serum proteins [50]. The substituents on the π-conjugated backbone of the ligand also affect the Cu^II/I^ reduction potentials. Whilst Cu(btsc) complexes are stable when the metal is in the +2 oxidation state, the reduction of the metal to copper(I) increases the susceptibility of the metal to dissociate from the ligand and transfer to copper proteins with a high affinity for copper(I) [51]. In general, modifying the aliphatic substituents on the N4 amine does not significantly affect the redox potential (±0.05 V), however it does have a pronounced effect on biodistribution and cellular uptake [52].

Here, Cu^II^(atsm) (**Figure 1A**) was selected as a reference compound due to its unique ability to penetrate the human BBB, which is a relatively rare characteristic among the metal complexes [53], [54]. Hence, the library of compounds primarily focused on the derivatives of Cu^II^(atsm) that retained electron-donating methyl or ethyl functional groups as to maintain similar CuII/I reduction potentials as Cu^II^(atsm). Additionally, compounds with methyl substituents on the backbone of the ligand were derived from 1,2-butanedione and were given an abbreviation starting with ‘a’ (in ex. Cu<u>A</u>TSM) while the compounds with ethyl backbone were derived from 3,4-hexanedione and were given an abbreviation starting with ‘d’ (in ex. Cu<u>D</u>TSM (Cu**L^1^**), Cu<u>D</u>TSE (Cu**L^2^**)). The substituents added in the N^4^- position were selected to add differing hydrogen bond donors and acceptors (alcohol (Cu**L^3^**), ester (Cu**L^4^**, Cu**L^5^**) and ether/polyethylene glycol functional groups, Cu**L^8^**, Cu**L^9^**, Cu**L^10^**) as well as morpholino (Cu**L^6^**) and pyridyl (Cu**L^7^**) functional groups. A general objective was to probe the different compounds for improved solubility in aqueous mixtures without significantly compromising their cell membrane permeability.

### Novel metal compounds exert various levels of cytotoxicity in human vascular endothelial cells

Since the majority of the tested here compounds have not been previously examined in human cell models, we first utilised the human umbilical vein endothelial cells (HUVEC) to perform a preliminary assessment of compound toxicity. The endothelial phenotype of utilised cells was confirmed by the observation of characteristic cobblestone-like morphology and the expression of the known endothelial cell marker vascular endothelial (VE)-cadherin (**Figure 1B**). To identify the range of non-toxic concentrations of designed Cu- and Ni-compounds, a cytotoxicity screen was performed utilising the colourimetric MTT assay. HUVEC were treated with increasing concentrations (0.1 μM - 3.0 μM) of each compound, corresponding to the range of Cu(ATSM) concentrations used in various *in vitro* cell models [42], [48], [55], [56] and cell viability was assessed after 24 h incubation with the compounds. Interestingly, we observed an expected dose-dependent decrease in the viability of HUVEC when treated with the majority of Cu compounds (**Figure 1C**). However, a similar effect was not observed in analogous Ni compounds, suggesting that Cu overload rather than the (btsc) backbone itself was driving the observed toxicity of high doses of tested compounds in this model (**Figure 1C**). In addition, we did not detect significant cytotoxic effects of compounds Cu**L^6^**and Cu**L^7^** at either of the tested concentration, indicating good biological tolerability of compounds with morpholino and pyridyl functional groups. Vehicle-only controls that corresponded to the two highest metal compound concentrations tested were also included, and these controls demonstrated no effect of vehicle treatment on HUVEC viability (**Figure S1A**).

Since cytotoxic effects were observed at the lowest concentrations ranging from 0.5 μM to 1.5 μM for certain compounds (Cu(ATSM), Cu**L^2^**, Cu**L^3^**, Cu**L^4^**, Cu**L^8^**, Cu**L^9^**; **Figure 1C**), two lower concentrations specifically 0.5 μM and 1.0 μM were selected for further testing of their effects on iBEC viability. Notably, a concentration of 0.1 μM was excluded from further analysis as it is below the robust detection limit of the inductively coupled plasma mass spectrometry (ICP-MS), which was used in subsequent experiments to measure Cu and Ni concentrations.

### Familial AD patient-derived iBEC serve as a novel approach for disease-specific metal compound screening

We next ulitised a familial AD patient-derived Transwell-based BBB model previously established and characterised by our group [40], [41], which involved the differentiation of iBEC from hiPSC lines of two AD patients carrying a disease-associated mutation (exon 9 deletion) in the presenilin-1 gene (*PSEN1-ΔE9*) [57] (**Figure 2A**, **Table 1**). Additional hiPSC lines were included as controls: one from an unrelated healthy donor line and two isogenic control lines where *PSEN1-ΔE9* mutation has been corrected with CRISPR-Cas9, as previously established by us [40], [57].

**Figure 2.**
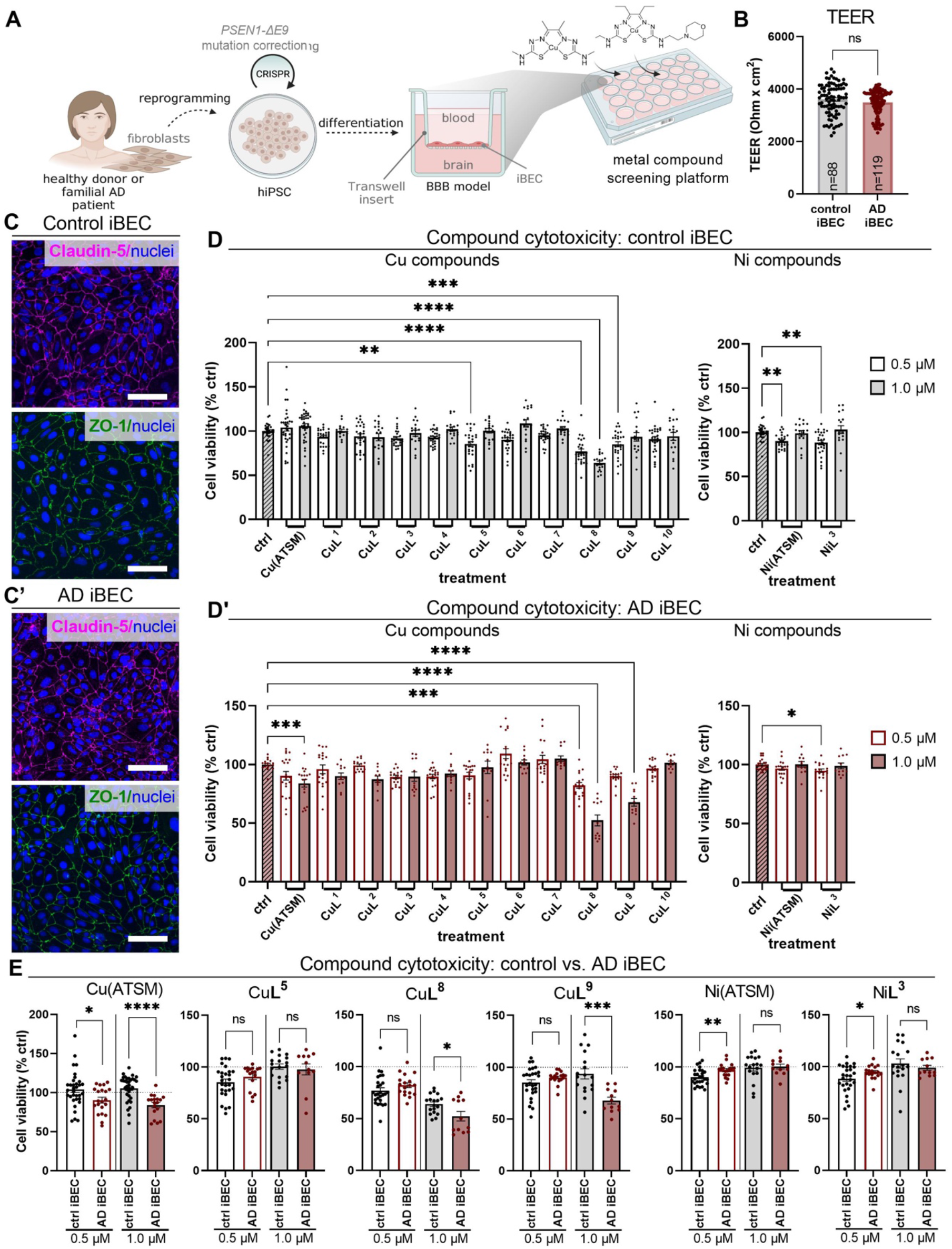
Effects of metal compounds on the viability of control and AD iBEC. (**A**) Schematic of established patient-derived metal compound screening platform. hiPSC carrying *PSEN1* mutation and respective control lines were differentiated towards brain endothelial-like cell phenotype and cultured in a Transwell insert to form a simple BBB *in vitro* model utilised in the metal compound screening assays. (**B**) Transendothelial electrical resistance (TEER) of monolayers formed by control and AD iBEC showed as Ohm x cm^2^. Control iBEC: N=3 lines, AD iBEC: N=2 lines; the number of independent replicates per cell group is indicated in the graph. (**C**) Representative immunofluorescence images of claudin-5 (magenta) and ZO-1 (green) with Hoechst nuclear counterstaining in control and AD iBEC. Scale bar, 100 μm. (**D- D’**) Control and AD iBEC viability after treatment with 0.5 μM and 1.0 μM of Cu and Ni compounds as assessed with MTT assay. Cell viability is shown as % of viable cells compared to untreated control (ctrl). (**E**) Comparison of the cytotoxic effects of selected metal compounds between control and AD iBEC. Cell viability is shown as % of viable cells as compared to respective untreated control, and compared between control and AD iBEC. In (D-E): Control iBEC: N=3 lines, AD iBEC: N=2 lines; n=8-9 independent replicates per line for 0.5 μM and n=5-6 independent replicates per line for 1.0 μM. Data are presented as mean ± SEM. Statistical analysis was performed using unpaired Welch’s t-test in (B,E) and one-way ANOVA with Dunnett’s test in (D-D’). \**p*<0.05, \*\**p*<0.01, \*\*\**p*<0.001, \*\*\*\**p*<0.0001. The dashed line represents untreated control.

**Table 1.** Summary of the controls and AD hiPSC lines included in the current study.

| Line ID | Age at biopsy | *PSEN1* genotype | *APOE* genotype |
| --- | --- | --- | --- |
| HDFa<br>(unrelated<br>healthy donor) | Not known | Not known | Not known |
| AD4 1.6.12.9 | 48 | *PSEN1-ΔE9* isogenic corrected | E3/E3 |
| AD5 1.5.6.1 | 47 | *PSEN1-ΔE9* isogenic corrected | E3/E3 |
| AD4 1.6 | 48 | *PSEN1-ΔE9* | E3/E3 |
| AD5 1.5 | 47 | *PSEN1-ΔE9* | E3/E3 |

To confirm the pluripotency of hiPSC, cells from all studied lines were characterised for the nuclear expression of stem cell-specific pluripotency markers including homeobox protein Nanog and (sex determining region Y)-box 2, known as Sox2, by immunofluorescence (**Figure S2**). Control and AD iBEC were generated from respective hiPSC lines using a previously published protocol [40], [58], [59](**Figure S3A-D**) and we confirmed the successful differentiation of hiPSC towards brain endothelial-like cell phenotype in this study through the expression of characteristic marker proteins: claudin-5, zonula occludens-1 (ZO-1), occludin, glucose transporter type 1 (Glut-1), and the formation of cobblestone-like confluent monolayer (**Figure 2C-C’**, **Figure S4**).

To further validate correct differentiation of control and AD iBEC towards the brain endothelial-like cell phenotype, we compared the expression of pluripotency genes *SOX2, NANOG* and octamer-binding transcription factor 4 *(OCT4),* and junctional and endothelial cell marker genes encoding for VE-cadherin (*CDH5*), claudin-5 (*CLDN5*), occludin (*OCLN*) and ZO-1 (*TJP1*) between undifferentiated hiPSC and iBEC generated from them. As anticipated, the expression of stem cell markers *SOX2*, *NANOG* and *OCT4* [60], [61], [62], were significantly higher in both control (*p*<0.0001) and AD (*p*<0.001) parental hiPSC when compared to iBEC (**Figure S3E**). These results confirmed the pluripotent nature of undifferentiated hiPSC lines and indicated that the expression of pluripotency genes becomes silenced upon lineage-specific differentiation in iBEC. Consistently, compared with undifferentiated hiPSC, iBEC generated from control and AD lines expressed increased levels of BBB and endothelial cell marker genes *CDH5* (control and AD lines: *p*<0.0001), *CLDN5* (control: *p*<0.001, AD: *p*<0.0001), *OCLN* (control and AD: *p*<0.001) and *TJP1* (control: *p*<0.001, AD: *p*<0.01), indicating their effective lineage-commitment towards a brain-endothelial cell-like phenotype (**Figure S3E’**).

Finally, to test for functional barrier formation in our model, control and AD iBEC were cultured in a Transwell insert and transendothelial electrical resistance (TEER) of the cell monolayer was measured with EVOM Volt/Ohmmeter. Both control and AD iBEC formed barriers with high TEER values (control iBEC: 3602 ± 63, AD iBEC: 3491 ± 44 Ohm x cm^2^, mean ± SEM) corresponding to the previously reported TEER range *in vivo* (1000 - 5900 Ohm x cm^2^, [63]– [65])(**Figure 2B**). No significant difference in barrier integrity was observed between control and AD iBEC (**Figure 2B**).

This established and characterised model was subsequently utilised as a patient cell-derived BBB platform for metal compound screening (**Figure 2A**).

### AD patient-derived iBEC demonstrate differential sensitivity to metal compound toxicity compared to control iBEC

With human BBB toxicity now emerging as an important concern in clinical trials [66], we first examined the effects of metal compound treatment on iBEC viability (at two pre-selected concentrations: 0.5 μM and 1.0 μM, for 24 h), and compared the responses between the control and AD cells.

The viability of control iBEC was significantly decreased after treatment with compounds Cu**L^5^** (*p*<0.01), Cu**L^8^** (*p*<0.0001), Cu**L^9^** (*p*<0.001) at 0.5 μM and compound Cu**L^8^** (*p*<0.0001) at 1.0 μM, as compared to untreated control (**Figure 2D**), revealing interesting inverse correlation of applied Cu compound concentration and control iBEC viability. A similar effect was observed for Ni compounds where Ni(ATSM) and Ni**L^3^** exerted significant (*p*<0.01) cytotoxic effects at 0.5 μM while not at 1.0 μM. The remaining compounds, as well as vehicle-only treatment, had no effect on control iBEC viability at two concentrations tested (**Figure 2D**, **Figure S1B**). When assessed in AD iBEC, compound Ni**L^3^** exhibited cytotoxicity at 0.5 μM (*p*<0.5), while Cu(ATSM) and Cu**L^9^** at 1.0 μM (*p*<0.001 and *p*<0.0001, respectively) (**Figure 2D’**). Similarly to control iBEC, treatment with compound Cu**L^8^** significantly decreased AD iBEC viability at both tested concentrations (0.5 μM: *p*<0.001, 1.0 μM: *p*<0.0001), confirming its high toxicity in human cell models. Intriguingly, classical dose-dependent toxicity was observed in AD iBEC for Cu compounds while the opposite trend was observed for Ni compound Ni**L^3^** which decreased cell viability only at the lower concentration tested (**Figure 2D’**). Other compounds as well as vehicle-only treatment were well tolerated by AD iBEC (**Figure 2D’**, **Figure S1B**).

We then focused on the group of compounds that significantly reduced cell viability in our BBB model (Cu(ATSM), Cu**L^5^**, Cu**L^8^**, Cu**L^9^**, Ni(ATSM) and Ni**L^3^**), and compared their effects between control and AD iBEC. When comparing within each tested concentration, AD iBEC proved to be more sensitive to Cu compounds Cu(ATSM) (*p*<0.0001), Cu**L^8^** (*p*<0.05) and Cu**L^9^** (*p*<0.001) treatment at 1.0 μM and Cu(ATSM) (*p*<0.05) at 0.5 μM, compared to control iBEC (**Figure 2E**). Interestingly, the opposite trend was observed for Ni compounds that had a stronger negative effect on control iBEC viability as compared to AD iBEC when tested at 0.5 μM (Ni(ATSM): *p*<0.01; Ni**L^3^**: *p*<0.05, **Figure 2E**). Among compounds that had no detrimental effect on cell viability, AD iBEC consistently presented a trend towards higher sensitivity to Cu compounds applied at 1.0 μM as compared to control iBEC for compounds Cu**L^2^** and Cu**L^3^**, with the effect being statistically significant for Cu**L^1^** (*p*<0.01) and Cu**L^4^** (*p*<0.01) (**Figure S5A**). Consistent with the results observed in HUVEC (**Figure 1C**), compounds Cu**L^6^** and Cu**L^7^** were not cytotoxic at either tested concentration, while at 0.5 μM these compounds appeared to improve the viability of AD iBEC when compared to control cells (Cu**L^6^**: control iBEC: 90.24 ± 2.117 vs AD: 109.3 ± 4.22; Cu**L^7^**: control iBEC: 95.09 ± 1.637 vs AD: 104.4 ± 3.416 % viability of untreated control, mean ± SEM, **Figure S5A**). This confirmed the lack of toxicity of these compounds in human cell models at tested concentrations and suggested their potential promising tolerability in AD patients. Finally, we observed differences in cell viability when directly comparing the effects of structurally analogous Cu and Ni compounds, Cu(ATSM) and Ni(ATSM), in control and AD iBEC, demonstrating opposite effect trends (**Figure S5B**). Namely, when comparing Cu**L^3^** vs Ni**L^3^** compound pair, we did not detect differential effects in control iBEC viability to those compounds, while AD iBEC were found to be more sensitive to Cu**L^3^** compared to Ni**L^3^** at 0.5 μM (*p*<0.05, **Figure S5B**).

Together these results provide a comprehensive characterisation of metal compound cytotoxicity in the human iPSC-derived BBB *in vitro* model, and identified differential sensitivity of AD iBEC to metal compound dose and chemical structure, compared to control cells.

### Compounds CuL^6^ and CuL^1^ demonstrate increased cell-associated accumulation in control and AD iBEC

Next, to gain insight into the accumulation of compounds at the BBB, control and AD iBEC cultured in the Transwell insert were treated with tested compounds and the levels of the corresponding metal accumulated in the cell monolayer were measured by ICP-MS (**Figure 3A**). Since the different chemical structure of each compound was designed to potentially alter its bioavailability and passive permeability through the brain endothelial cell membrane, we sought to test all the compounds at the same concentration to enable comparison of the effects of the (btsc) ligand backbone on the cellular uptake of the compound. Given that some of the tested compounds demonstrated various effects on cell viability depending on the iBEC genotype (**Figure 2D-D’**), we selected 0.5 μM concentration for further investigation. This concentration was better tolerated by AD iBEC (**Figure 2D’**), and aligns with our primary interest in AD cells, offering higher clinical relevance. However, to minimise the confounding effect of compound cytotoxicity in iBEC, we shortened the compound incubation time with iBEC from 24 h to 2 h hypothesizing that this shorter exposure period would incur less pronounced effects on cell viability.

**Figure 3.**
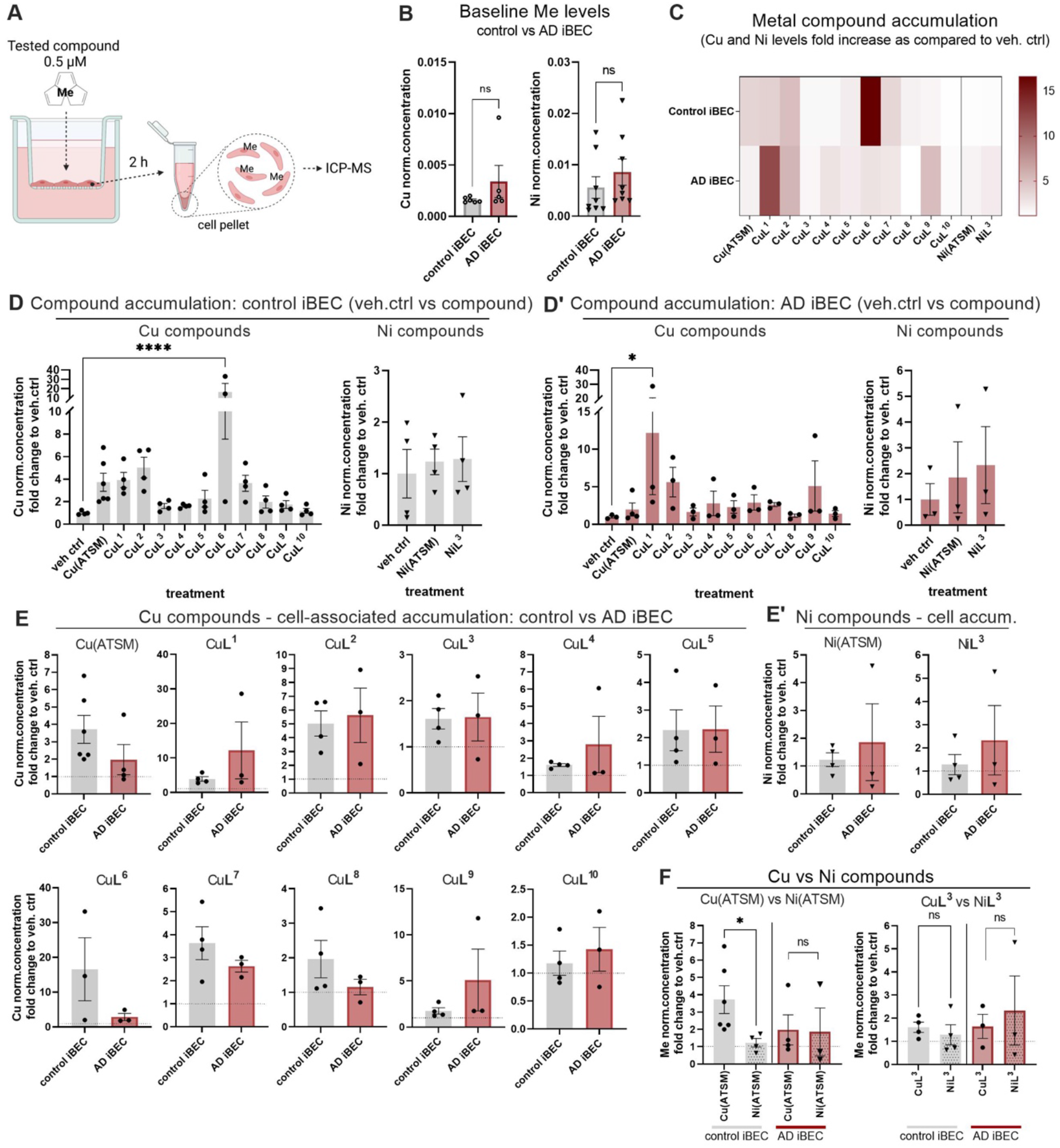
Cell-associated accumulation of tested compounds in control and AD iBEC. (**A**) Schematic of experimental workflow. To determine the cell-associated accumulation of tested compounds, iBEC derived from control and AD hiPSC were cultured on Transwell inserts and compounds were added at 0.5 μM to the top chamber of the Transwell insert. Following 2 h treatment, cell pellet samples were collected and Cu and Ni concentrations [μmol/L] were assessed with ICP-MS. (**B**) Comparison of normalised Cu and Ni levels in experimental control (untreated and vehicle-only treated) control and AD iBEC. Cu and Ni levels were normalised to Mg concentration in every individual sample and compared between control and AD iBEC. (Control iBEC: N=3 lines, AD iBEC: N=2 lines; n=1-3 independent replicates per line). (**C**) Heatmap summarising fold changes in the cell-associated metal accumulation in control and AD iBEC. (**D-D’**) The accumulation of cell-associated Cu and Ni in control and AD iBEC as compared to vehicle-treated control. (**E-E’**) Comparison of Cu and Ni accumulation in control vs AD iBEC. (**F**) Comparison of metal accumulation between Cu and Ni compounds in control and AD iBEC. In (C-F) Cu or Ni levels were normalised to Mg concentration in every individual sample and presented as a fold change of Cu and Ni normalised (norm.) levels as compared to the respective vehicle-treated control. (Control iBEC: N=3 lines, AD iBEC: N=2 lines; n=1-2 independent replicates per line). Statistical analysis was performed using unpaired Welch’s t-test in (B, E-F) and one-way ANOVA with Dunnett’s test in (D-D’). \**p*<0.05, \*\**p*<0.01, \*\*\**p*<0.001, \*\*\*\**p*<0.0001. Me-metal. The dashed line represents vehicle-treated control.

Since AD was previously associated with the dysregulation of Cu homeostasis in the brain [67], we first compared the baseline levels of cell-associated Cu between control and AD iBEC (**Figure 3B**). In the absence of metal compound treatment, we observed a small trend toward increased levels of Cu in AD iBEC as compared to control cells, suggesting that this phenotype is not strongly present in the AD patient-derived iBEC.

In our metal compound treatment experiment, we observed trends indicating a higher level of cell-associated metal in both control and AD iBEC as compared to vehicle-only controls (**Figure 3C-D’**). This suggests that the tested metal-(btsc)s are cell membrane-permeable and can accumulate intracellularly *in vitro*. Interestingly, when compared to vehicle controls, the levels of Cu were significantly increased in control iBEC treated with compound Cu**L^6^**(16.61 ± 9.05 fold increase in Cu *vs*. vehicle ctrl, mean ± SEM, *p*<0.0001, **Figure 3C-D**), while Cu**L^1^**demonstrated increased accumulation in AD iBEC (12.18 ± 8.24 fold increase in Cu *vs*. vehicle ctrl, mean ± SEM, *p*<0.05, **Figure 3C and D’**), suggesting their potentially improved cellular uptake by BBB cells. The limitation of this experiment however was substantial variability among independent replicates, with compounds Cu**L^2^**and Cu**L^7^** demonstrating more consistent, although moderate elevation of Cu levels in iBEC (**Figure 3D-D’**).

In order to identify a potential drug candidate with superior delivery specifically in the context of AD, we compared the efficiency of compound accumulation between control and AD iBEC treated with tested (btsc)s (**Figure 3E-E’**). We found variable effects where Cu(ATSM), Cu**L^6^**, Cu**L^7^**and Cu**L^8^**, showed a trend towards lower uptake in AD iBEC compared to control cells while Cu**L^2^**, Cu**L^3^**, Cu**L^5^** and Cu**L^10^** demonstrated a similar level of cellular accumulation between control and AD iBEC. While only trends were observed, we identified compounds Cu**L^1^**, Cu**L^4^**, and Cu**L^9^**, as complexes with potentially better accumulation in AD cells compared to control iBEC (**Figure 3E**).

Interestingly, when comparing the efficiency of cellular accumulation between Cu and Ni structural analogues, we observed an increase in metal accumulation delivered by Cu(ATSM) as compared to Ni(ATSM) in control cells (*p*<0.05) (**Figure 3F**). However, no differences were observed in AD cells or between Cu**L^3^** and Ni**L^3^** compound pair (**Figure 3F**). These findings highlight the complex relationship between the ligand backbone and the central metal of the compound as well as the genetic background of the cells in terms of transport at the human BBB level.

### Compound permeability screen identifies improved delivery of Cu(ATSM) and CuL^1^ in AD patient-derived BBB

To further compare the penetration of metal compounds across the human BBB *in vitro*, control and AD iBEC were cultured on Transwell inserts to form a tight cell monolayer and metal compounds were added at a concentration of 0.5 μM to the top chamber of the Transwell insert. Following 2 h incubation, the cell culture media from the bottom chamber of the Transwell insert was collected for assessment of Cu and Ni concentrations using ICP-MS (**Figure 4A**).

**Figure 4.**
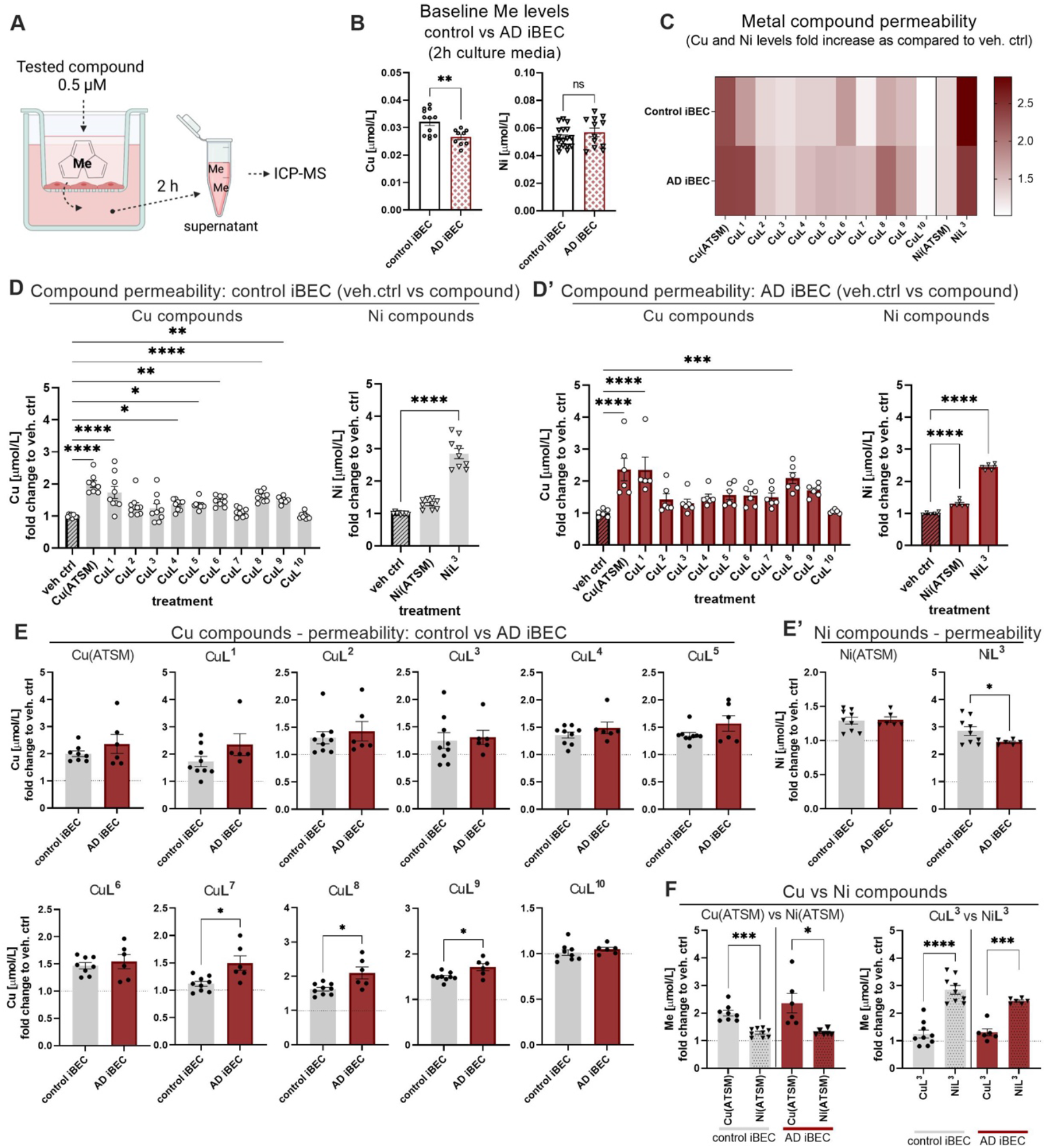
Permeability of metals through the iBEC monolayer after treatment with tested compounds. (**A**) Schematic of experimental workflow. To determine the permeability of metal delivered by tested compounds in the BBB model, iBEC derived from control and AD hiPSC were cultured on Transwell inserts and compounds were added at 0.5 μM to the top chamber of the Transwell insert. Following 2 h treatment, media samples were collected from the bottom chamber of the Transwell insert and Cu and Ni concentrations were assessed with ICP-MS. (**B**) Comparison of Cu and Ni concentration [μmol/L] measured in cell culture media collected from wells corresponding to ‘pooled control’ (untreated and vehicle-only treated) control and AD iBEC, at 2 h post media change (Control iBEC: N=3 lines, AD iBEC: N=2 lines; a minimum of n=3 independent replicates per line). (**C**) Heatmap summarising fold changes in the permeability of metal compounds through iBEC monolayer formed by control and AD cells. (**D-D’**) The permeability of Cu and Ni delivered by tested compounds in control and AD iBEC as compared to vehicle-treated control. (**E-E’**) Comparison of metal permeability from compound treatment in control vs AD iBEC. (**F**) Comparison of the efficacy of metal permeability between Cu and Ni compounds in control and AD iBEC. Results in (C-F) are presented as a fold change of Cu and Ni concentration [µmol/L] in the cell media collected from the bottom chamber of Transwell insert in metal compound-treated well, as compared to the respective vehicle-treated control. (Control iBEC: N=3 lines, AD iBEC: N=2 lines; n=2-3 independent replicates per line). Statistical analysis was performed using unpaired Welch’s t-test in (B, E-F) and one-way ANOVA with Dunnett’s test in (D-D’). \**p*<0.05, \*\**p*<0.01, \*\*\**p*<0.001, \*\*\*\**p*<0.0001. Me-metal. The dashed line represents vehicle-treated control.

To understand the baseline levels of Cu and Ni in our culture conditions, we first compared the concentrations of these metals in cell culture media collected from experimental control (untreated and vehicle-only treated) Transwells from control and AD iBEC after 2 h. Results revealed a similar level of Ni in media collected from wells where control or AD iBEC were cultured, however, we detected lower (*p*<0.01) concentrations of Cu in media collected from wells corresponding to AD iBEC as opposed to those with control iBEC (**Figure 4B**). Additionally, no differences in Cu or Ni levels were detected between untreated and vehicle-only treated cells (data not shown). As we observed a small trend towards increased levels of Cu in AD iBEC cell pellets collected from the corresponding Transwell inserts (**Figure 3B**), this suggests that AD cells may regulate Cu differently compared to control iBEC. Importantly, the baseline Cu levels detected in cell culture media were at least 15-fold lower than our selected Cu compound treatment concentration (0.5 μmol/L) (control iBEC: 0.0321 ± 0.001 μmol/L Cu, AD iBEC: 0.0265 ± 0.001 μmol/L Cu, mean ± SEM, **Figure 4B)**. Therefore, the differences in baseline Cu levels were unlikely to confound our assessment of Cu concentration in the Transwell flow-through media samples collected during the compound permeability experiments.

Correspondingly, in our metal compound permeability screen, we observed trends toward increased concentration of Cu and Ni in the media collected from compound-treated wells as compared to vehicle-only controls, suggesting effective transport of metal-(btsc)s, or associated metal, through iBEC monolayers (**Figure 4C-D’**). This transport may occur *via* simple gradient-driven diffusion, although the involvement of active transport cannot be ruled out [43], [68], [69]. Interestingly, treatment with several compounds, including Cu**L^1^** (*p*<0.0001), Cu**L^4^** (*p*<0.05), Cu**L^5^**(*p*<0.05), Cu**L^6^** (*p*<0.01), Cu**L^9^** (*p*<0.01) as well as Ni**L^3^** (*p*<0.0001) resulted in higher Cu and Ni levels, respectively, in the media samples collected from the compound-treated wells (as compared to vehicle-only controls) in control iBEC, suggesting human *in vitro* BBB permeability (**Figure 4C-D**). Notably, Cu compound Cu**L^1^** demonstrated increased permeability in AD iBEC (Cu**L^1^**: 2.35 ± 0.4 fold increase in Cu [μmol/L] *vs*. vehicle ctrl, mean ± SEM, *p*<0.0001; **Figure 4C, D’**), while most of the novel Cu compounds achieved only moderate permeability in AD cells (**Figure 4D’**). Cu(ATSM), serving here as a positive control, demonstrated increased permeability in both control (*p*<0.0001) and AD (*p*<0.0001) iBEC monolayers (**Figure 4D-D’**), confirming the relevance of our model to the human BBB [53], [54]. Both Ni compounds also showed increased (*p*<0.0001) permeability in AD iBEC (**Figure 4D’**).

Compound Cu**L^10^** consistently demonstrated poor permeability in control and AD iBEC monolayer, presenting limited translational potential in the context of brain disorders. Additionally, the permeability of Cu**L^8^** was significantly increased in control (*p*<0.0001) and AD iBEC (*p*<0.001). However, this effect is likely attributed to its high cytotoxicity (**Figure 1C**, **Figure 2D-D**’) and therefore potential undesired disruption of the iBEC monolayer.

When compared between control and AD iBEC, the permeability efficiency of tested compounds largely did not differ between the two cell groups, except for Cu**L^7^**, Cu**L^8^** and Cu**L^9^**which showed higher (*p*<0.5) permeability in AD iBEC (**Figure 4E-E’**). Contrarily, the permeability efficiency of Ni**L^3^** was significantly lower (*p*<0.05) in AD iBEC as compared to control iBEC (**Figure 4E’**). Interestingly, we did not find significant differences in the iBEC monolayer integrity between control and AD iBEC (**Figure 2B**) suggesting that the observed differences in metal compound passage through the BBB *in vitro* can be primarily attributed to differences in the chemical structure of tested Cu- and Ni-(btsc). Intriguingly, when comparing structurally analogous Cu and Ni compounds, we observed higher permeability efficacy of Cu(ATSM) as compared to Ni(ATSM), while the opposite was found for Cu**L^3^** and Ni**L^3^**compound pair where Ni**L^3^** showed significantly increased permeability compared to its Cu structural analogue (**Figure 4F**). Together these results highlight that the differential effects of compound permeability may be driven by both the conjugated central metal and the modifications applied to (btsc) ligand.

### AD patient-derived iBEC exhibit an inflammatory response to TNFα and IFNγ stimulation

Since our previous studies demonstrated anti-neuroinflammatory effects of Cu(btsc) complexes in various models of neurodegeneration [42], [48], [49], here we hypothesised that novel metal (btsc) complexes can achieve similar effects in iBEC and aimed to evaluate the immunomodulatory activity of selected compounds in the established AD-patient-derived platform.

Although hiPSC-derived iBEC were previously shown to respond to proinflammatory mediators such as TNFα [35], [70], [71], a comprehensive characterisation of their immunophenotype in the context of familial AD has not yet been described.

We therefore first evaluated the baseline inflammatory profile of cells in our BBB models *via* qPCR. Interestingly, there were no significant differences in the gene expression of classical proinflammatory markers including interleukin-6 (*IL6*), monocyte chemoattractant protein-1 (*CCL2*), interleukin-8 (*IL8*), interleukin-1α (*IL1A*) or tumour necrosis factor-α (*TNF*) between control and AD iBEC (**Figure S6A**), while the interleukin-1β (*IL1B*) was below detection levels. Under baseline conditions, the expression of oxidative stress markers also was not altered in AD iBEC compared to control cells (**Figure S6B**). Collectively, these results suggest a minimal inflammatory response in AD patient-derived iBEC under normal conditions, which contrasts with previous observations for BEC in the human AD brain [72]–[74]. This discrepancy may be due to the specific platform and culture conditions employed for AD iBEC. To address this, we aimed to mimic the disease-associated inflammatory microenvironment observed in the AD brain by stimulating AD patient-derived iBEC with neurologically relevant proinflammatory mediators TNFα and IFNγ [2], [75]–[77] for 24 h before conducting a range of assays (**Figure 5A**). Utilised concentrations of TNFα (20 ng/ml) and IFNγ (30 ng/ml) were selected based on established literature [78]–[82].

**Figure 5.**
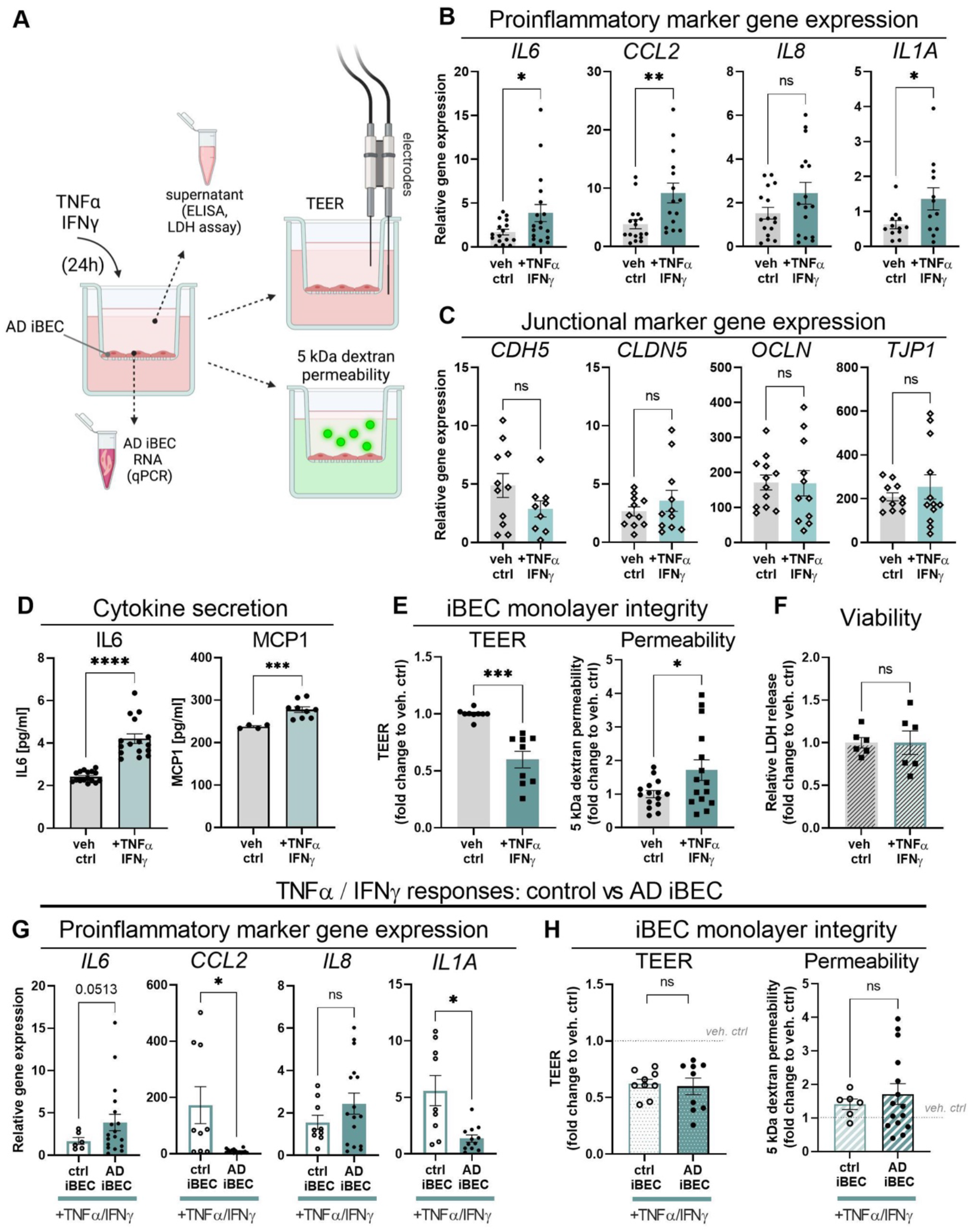
Effects of TNFα/IFNγ- stimulation on AD iBEC phenotype. (**A**) Schematic of experimental workflow. To determine the effects of TNFα/IFNγ- stimulation on the AD patient-derived BBB model, AD iBEC were cultured on Transwell inserts and TNFα/IFNγ were added to the top chamber of the Transwell insert. Following 24 h treatment, TEER measurement and 5 kDa dextran permeability assays were performed. Cell pellet and supernatant samples were collected for subsequent analysis with qPCR, ELISA and LDH assays. (**B**) Relative expression of mRNA for proinflammatory marker genes *IL6*, *CCL2*, *IL8* and *IL1A* in vehicle- and TNFα/IFNγ- treated AD iBEC. Results presented as ΔΔCT x 10^6^. (AD iBEC: N=2 lines, minimum n=5 independent replicates per line). (**C**) Relative expression of mRNA for tight and adherens junctional marker genes *CDH5*, *CLDN5*, *OCLN* and *TJP1* in vehicle- and TNFα/IFNγ- treated AD iBEC. Results presented as ΔΔCT x 10^6^. (AD iBEC: N=2 lines, minimum n=3 independent replicates per line). (**D**) Secretion of proinflammatory cytokines IL6 and MCP1 in vehicle-treated and TNFα/IFN-treated AD iBEC. Data showed as [pg/ml] concentration of each cytokine in cell supernatant at 24 h post-treatment (AD iBEC: N=2 lines, IL6: minimum n=4 independent replicates per line; MCP1: minimum n=2 independent replicates per line. Additional n=5 vehicle-treated control samples were analysed for MCP1 levels. The resulting normalised absorbance values were 0, -0.001 or -0.0005 following background and blank subtraction suggesting MCP1 levels being below the detection limit of the assay, and therefore excluded from analysis.) (**E**) Changes in AD iBEC monolayer TEER and passive permeability to 5 kDa dextran following treatment with TNFα/IFNγ. Left panel: data showed as fold change in TEER as compared to vehicle-treated control at 24 h (AD iBEC: N=2 lines, minimum n=3 independent replicates per line). Right panel: data showed as fold change in 5 kDa dextran clearance volume to vehicle-treated control at 24 h (AD iBEC: N=2 lines, minimum n=5 independent replicates per line). (**F**) Relative lactate dehydrogenase (LDH) release in AD iBEC after stimulation with TNFα/IFNγ. LDH release showed as fold changes to vehicle-treated control (AD iBEC: N=2 lines, n=3 independent replicates per line). (**G**) Comparison of relative expression of mRNA for proinflammatory marker genes *IL6*, *CCL2*, *IL8* and *IL1A* in vehicle- and TNFα/IFNγ- treated control and AD iBEC. Results presented as ΔΔCT x 10^6^. (Control iBEC: N=3 lines, AD iBEC: N=2 lines, n=1-3 independent replicates per line). (**H**) Comparison of changes in control iBEC and AD iBEC monolayer integrity following treatment with TNFα/IFNγ. Left panel: monolayer TEER. Data showed as fold change in TEER as compared to the respective vehicle-treated control at 24 h. Right panel: monolayer passive permeability. Data showed as fold change in 5 kDa dextran clearance volume to respective vehicle-treated control at 24 h (Control iBEC: N=2 lines, AD iBEC: N=2 lines, minimum n=3 independent replicates per line). Statistical analysis was performed using unpaired Welch’s t-test in (B-H). \**p*<0.05, \*\**p*<0.01, \*\*\**p*<0.001, \*\*\*\**p*<0.0001. veh ctrl- vehicle-treated control;

In TNFα/IFNγ-treated AD iBEC, we observed an increased expression of *IL6* (*p*<0.05), *CCL2* (*p*<0.01) and *IL1A* (*p*<0.05) compared to vehicle-treated control (**Figure 5B**). Additionally, there was a trend towards increased *IL8* expression following TNFα/IFNγ exposure (veh ctrl: 1.53 ± 0.27, TNFα/IFNγ: 2.44 ± 0.5 relative *IL8* gene expression, mean ± SEM, *p*=0.1204; **Figure 5B**). Furthermore, our gene expression results corroborate with the corresponding protein secretion profiles where a significant increase in the secretion of IL6 (*p*<0.0001) and MCP1 (*p*<0.001) was detected in TNFα/IFNγ-treated AD iBEC (**Figure 5D**). This confirms the activation of pro-inflammatory pathways in AD iBEC in response to TNFα/IFNγ stimulation at both the transcriptional and protein level.

TNFα and interferons were previously reported to decrease tight and adherent junction expression and affect the functional characteristics of human BEC [35], [83]. Therefore, we next assessed the impact of TNFα/IFNγ on the integrity of BBB in our AD model. While TNFα/IFNγ treatment did not elicit significant changes in the expression of junctional markers *CDH5*, *CLDN5*, *OCLN* and *TJP1* (**Figure 5C**), potentially due to already reduced baseline expression of those genes in AD iBEC (**Figure S6C**), it resulted in decreased TEER (*p*<0.001) and increased passive permeability to biologically inert fluorescent tracer (5 kDa FITC- conjugated dextran, *p*<0.05) in AD iBEC monolayers following exposure to TNFα/IFNγ (**Figure 5E**). These findings are consistent with the functional impairment of BBB integrity reported during AD-related neuroinflammation (**Figure 5E**) [25].

Interestingly, TNFα/IFNγ treatment induced a similar phenotype in control iBEC (**Figure S7A- B**) but the expression profile of proinflammatory markers differed between control and AD iBEC (**Figure 5G**). Specifically, *CCL2* and *IL1A* were expressed at higher (*p*<0.05) levels in control iBEC as compared to AD iBEC following TNFα/IFNγ stimulation, while *IL6* and *IL8* showed a trend towards higher expression in AD cells as compared to controls treated with TNFα/IFNγ (**Figure 5G**). Moreover, no significant differences in TNFα/IFNγ-induced changes in monolayer integrity were observed between the control and AD iBEC (**Figure 5H**). These findings suggest that distinct immune pathways may contribute to similar functional impairments in control and AD iBEC. Importantly, the observed effects were not induced by adverse changes in cell viability of control or AD iBEC following TNFα/IFNγ exposure (**Figure S7C**, **Figure 5F**).

Together these results illustrate that AD patient-derived iBEC are responsive to cytokine activation by the development of a characteristic, AD-relevant [4], [25], [26] proinflammatory phenotype including iBEC monolayer integrity impairment. This demonstrates the functional capability of our patient-derived iBEC Transwell system to model BBB neuroinflammation in AD and provides a useful tool for the further investigation of the immunomodulatory effects of metal compounds within a single experimental platform.

### CuL^1^ reverses TNFα/IFNγ-induced neuroinflammatory phenotype in AD iBEC

Given that Cu(btsc) have previously demonstrated robust anti-neuroinflammatory actions in preclinical models of neurodegeneration (microglia, astrocytes and neuronal *in vitro* cultures, and animal models [42], [48], [49]), we anticipated compounds in our library could exert similar, potentially therapeutic effects. Based on compound chemistry, toxicity, cellular accumulation and iBEC monolayer permeability, we selected Cu(ATSM), its control Ni(ATSM), and a group of five promising Cu compounds (Cu**L^1^**, Cu**L^4^**, Cu**L^6^**, Cu**L^7^** and Cu**L^9^**) for their further assessment in our model. Considering the clinical relevance, we focused on AD patient-derived iBEC to identify potential therapeutic effects of these metal-(btsc)s.

To assess the immunomodulatory properties of metal compounds, AD iBEC were treated with TNFα/IFNγ alone or in co-treatment with selected compound at 0.5 μM for 24 h. The responses of cells were examined across a panel of various established assays (**Figure 5A**). Importantly, none of the selected compounds showed cytotoxicity at this concentration and treatment duration in AD iBEC (**Figure 2D’**).

Interestingly, at the gene expression level, TNFα/IFNγ co-treatment with selected Cu compounds led to a significant decrease of at least one of the tested (*IL6*, *IL8*, *IL1A*, *CCL2*) proinflammatory marker genes as compared to TNFα/IFNγ alone (**Figure 6A**), supporting previously reported anti-inflammatory properties of Cu(btsc) [42], [48], [49]. Compound Cu**L^1^**emerged as the most promising candidate that, in co-treatment with TNFα/IFNγ, reduced expression of all four marker genes tested (*IL6*: *p*<0.05, *CCL2*: *p*<0.001, *IL8*: *p*<0.05, *IL1A*: *p*<0.001), as compared to TNFα/IFNγ alone (**Figure 6A**). Intriguingly, a decrease in gene expression of proinflammatory markers did not result in changes in cytokine secretion as the levels of IL6 and MCP1 were similar between TNFα/IFNγ and TNFα/IFNγ+metal compound treated cells at tested 24 h timepoint (**Figure 6B**). It has been shown however that BEC generate a temporarily dynamic cytokine secretion profile following activation with proinflammatory mediators [82], and therefore it is possible we did not capture those changes at a single time point tested. Similarly, others have found cytokine mRNA and protein production to peak at defined time points following cell activation [84]–[86], further suggesting that cytokine gene expression changes and resulting protein synthesis follow distinct kinetics in our model and applied treatment, and may not be detectable when assessed at single timepoint.

**Figure 6.**
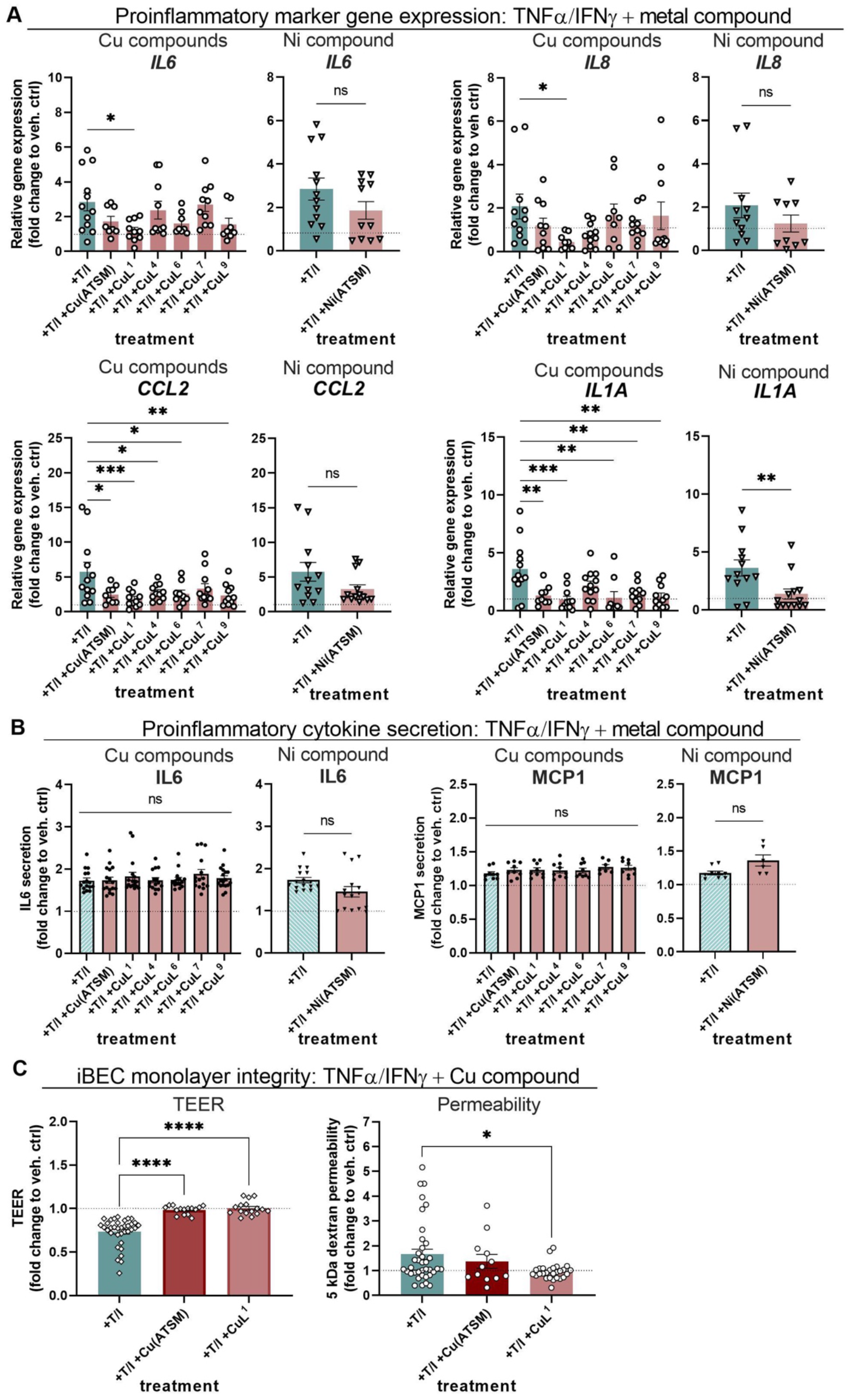
Anti-inflammatory effects of metal compounds in AD iBEC. (**A**) Relative expression of mRNA for *IL6*, *CCL2, IL8*, and *IL1A* in AD iBEC after stimulation with TNFα/IFNγ alone or together with 0.5 μM of tested compounds for 24 h. Results presented as fold change in ΔΔCT x 10^6^ as compared to vehicle-treated control (AD iBEC: N=2 lines, minimum n=3 independent replicates per line). (**B**) Secretion of proinflammatory cytokines IL6 and MCP1 in AD iBEC after stimulation with TNFα/IFNγ alone or together with 0.5 μM of tested compounds for 24 h. Data showed as [pg/ml] concentration of each cytokine in cell supernatant at 24 h post treatment. (AD iBEC: N=2 lines, minimum n=3 independent replicates per line). (**C**) Changes in AD iBEC monolayer TEER and passive permeability to 5 kDa dextran following 24 h treatment with TNFα/IFNγ alone or in combination with 0.5 μM Cu(ATSM) or Cu**L^1^**. Left panel: data showed as fold change in TEER as compared to vehicle-treated control at 24 h (AD iBEC: N=2 lines, minimum n=6 independent replicates per line). Right panel: data showed as fold change in 5 kDa dextran clearance volume to vehicle-treated control at 24 h (AD iBEC: N=2 lines, minimum n=5 independent replicates per line). Statistical analysis in (A-C) was performed using one-way ANOVA with Dunnett’s test for Cu compounds and unpaired Welch’s t-test for Ni compounds. \**p*<0.05, \*\**p*<0.01, \*\*\**p*<0.001, \*\*\*\**p*<0.0001. veh. ctrl- vehicle-treated control; T/I- TNFα and IFNγ. The dashed line represents vehicle-treated control.

In contrast, Ni(ATSM), exhibited minimal anti-inflammatory effects in AD iBEC with a significant decrease observed only in the expression of *IL1A* (*p*<0.01)(**Figure 6A**). However, when compared to Cu(ATSM), the immunomodulatory responses elicited by Ni(ATSM) did not differ from its Cu analogue when co-treated with TNFα/IFNγ (**Figure S8A-B**). Similar observations have been previously reported for Cu(ATSM) and Ni(ATSM) effects on ferroptosis and lipid peroxidation in N2 cells and cell-free systems respectively [56], together highlighting the importance of the conjugated ligand backbone for metal-(btsc) biological activity.

Finally, to assess the functional changes induced by TNFα/IFNγ in the AD BBB, we focused on the compound Cu**L^1^**, the most promising candidate identified in our qPCR screen (**Figure 6A**). The anti-inflammatory properties of Cu(ATSM) have been validated by others and in our studies [42], [48], making it a relevant reference compound in this case.

Interestingly, Cu**L^1^** treatment effectively prevented the detrimental effects of TNFα/IFNγ on AD iBEC barrier integrity as evidenced by the significant improvement in TEER (*p*<0.0001) and normalised passive permeability to 5 kDa FITC-conjugated dextran (*p*<0.05) (**Figure 6C**). These findings provide the first evidence that Cu(btsc) can ameliorate AD-related neuroinflammatory changes in BEC phenotype and function. Additionally, co-treatment with Cu(ATSM) rescued changes in TEER induced by TNFα/IFNγ (*p*<0.0001), although it did not significantly affect monolayer permeability to 5 kDa dextran suggesting the selective beneficial effect of this compound in AD iBEC (**Figure 6C**). Importantly, treatment with Cu**L^1^** or Cu(ATSM) alone had no effect on AD iBEC TEER or passive permeability to 5 kDa dextran confirming the lack of intrinsic effects of these compounds on iBEC monolayer integrity and function (**Figure S8C-D**).

Together, the results outline the practical application of our neuroinflammation-like AD iBEC model and validate the use of an array of assays to evaluate the immunomodulatory effects of novel metal compounds *in vitro*. Importantly, our study and compound screen is the first to identify the beneficial anti-neuroinflammatory effects of Cu(btsc) specifically in BEC, offering a novel mechanistic insight into their therapeutic potential in AD neurodegeneration.

## DISCUSSION

Despite a large number of clinical trials performed, therapeutics tested so far have largely failed to demonstrate robust symptom improvement in AD patients and AD has no cure [87]– [90]. This calls for a paradigm shift in AD drug discovery that may originate from the development of AD patient-specific preclinical drug screening platforms and early identification of therapeutics that can be successfully delivered across the BBB.

Respectively, hiPSC-derived *in vitro* BBB models incorporating human iBEC have recently emerged as a promising drug candidate screening tool offering the highest human *in vivo* BBB permeability prediction accuracy [31], [32]. Although transcriptomic meta-analysis suggests that hiPSC-derived iBEC demonstrate a partial component of epithelial identity [91], [92], iBEC currently serve as a state-of-the-art cell source for human BBB modelling, specifically in a diseases-context, and offer a high predictive capacity of human BBB function in health and disease [93]. However, despite providing an attractive alternative to traditionally used Caco-2, MDCK, or PAMPA drug permeability assays, the practical utility of hiPSC-derived BBB models is yet to be tested in large-scale AD drug screening studies.

Here we propose a unique patient cell-derived human BBB model as a validated platform for screening of new metal-(btsc) compound delivery and anti-neuroinflammatory activity in a familial AD context. hiPSC-derived iBEC utilised in this model demonstrated physiologically relevant, *in vivo* BBB-like characteristics such as high barrier integrity and expression of relevant markers. Simultaneously, AD patient-specific cells were shown to be compatible with a panel of medium- to high-throughput drug candidate testing BBB assays, including an assessment of their anti-inflammatory effects. With hiPSC offering in theory an infinite source of patient-derived cells, this BBB model offers a highly scalable platform that can be routinely used in anti-neuroinflammatory drug screening. Since in the current clinical trials, compounds targeting inflammatory pathways compose the largest percentage of disease-modifying therapies being tested [94], we believe this model and compound library being highly relevant for ongoing anti-neuroinflammatory drug discovery efforts in AD.

### hiPSC-derived AD models may aid in predicting metal compound toxicity at patient’s BBB

Cu(btsc) are small molecule compounds with proven immunomodulatory effects in preclinical models of neurodegeneration [42], [48], [49]. Extensive evidence demonstrates that the chemical properties and biological activity of metal (btsc) complexes are remarkably sensitive to the substituents attached to the backbone of the ligand and even small modifications of the ligand framework results in dramatic changes to complex stability, membrane permeability, redox activity, lipophilicity, interactions with serum albumin, cellular metal ion bioavailability and finally, therapeutic activity [44], [51], [69], [95], [96]. Here, we designed a library of novel metal-(btsc) compounds incorporating various modifications to its ligand and trialled their resulting cytotoxicity, cellular accumulation and BBB permeability in an established AD patient-derived model.

When assessing the cytotoxicity of novel metal compounds in HUVEC and iBEC, we identified relatively high toxicity of compounds Cu**L^8^** and Cu**L^9^** as compared to other tested metal-(btsc)s indicating low tolerance of Cu(btsc) containing polyethylene glycol (PEG) chains in their ligand structure. Conversely, treatment with compounds Cu**L^6^** and Cu**L^7^** had no cytotoxic effects at either concentration tested in both HUVEC and iBEC models, but seemingly improved the viability of AD iBEC when applied at the lower dose, suggesting potential protective effects of Cu complexes harbouring morpholino (Cu**L^6^**) and pyridyl (Cu**L^7^**) functional groups in AD cells. We observed also differences in compound cytotoxicity when compared between different tested complexes, collectively demonstrating that slight modifications to both central metal and (btsc) ligand backbone modulate the compound’s effect on human endothelial cell viability. In line with our observation, similar results were reported for human immortalised endothelial cells that demonstrated differential sensitivity to Cu^II^(atsm) and Cu^II^(gtsm) *in vitro* [55].

Our experiments also revealed the differences in viability of HUVEC and iBEC treated with the same library of compounds applied at the matching concentration and treatment duration. In detail, except for Cu**L^8^**, none of the compounds that decreased viability in iBEC at 0.5 μM were cytotoxic in HUVEC at this concentration, while simultaneously, compounds Cu**L^2^**and Cu**L^4^**, which demonstrated significant negative effects on the viability of HUVEC at 1.0 μM, were well tolerated by control and AD iBEC at the same dose. This suggests important differences in the sensitivity of hiPSC-derived iBEC as opposed to primary and non-CNS endothelial cell lines, further highlighting the challenging results discrepancy between traditional and hiPSC-derived BBB models. With iBEC demonstrating more *in vivo* BBB-like phenotype and function as compared to HUVEC [22], one could speculate the metal compound toxicity observed here in the iBEC model is more predictive of the clinically relevant toxic dose for human patients, however, this remains yet to be elucidated.

We further discovered an interesting correlation between cell genotype and metal delivered by (btsc) complexes in regard to cytotoxicity, where AD iBEC demonstrated generally higher vulnerability to Cu compounds while control iBEC were more sensitive to tested Ni complexes.

That differential sensitivity of control and AD patient-derived cells to metal (btsc) highlights the importance of drug candidate toxicity testing in patient-specific models to more reliably predict compound effects in the patient brain *in vivo*. Consequently, with BBB cells being altered in AD [4], [19], patient cells may prove to be more vulnerable to particular drug cytotoxicity and therefore drug doses and formulations tested in clinical trials should be precisely adjusted to match patient-specific BEC molecular and functional profiles. With hiPSC depositories now offering a variety of patient-derived lines with clearly defined genetic profiles, *in vitro* models as presented in this study may serve as a valuable drug candidate response prediction tool to test for potential BBB toxicity in sub-cohorts of AD patients.

### Cellular accumulation and permeability assays identify compounds with improved delivery in patient-derived BBB model

With BBB serving as a key hurdle in successful CNS drug delivery in AD [5], [12], we next aimed to validate our Transwell-based patient-cell derived iBEC model for compound cellular accumulation and permeability assays.

Here were able to identify compound Cu**L^1^** as a complex of specifically high cellular uptake in AD iBEC, suggesting that certain chemical modifications to (btsc) framework, such as the ethyl backbone (as in Cu**L^1^**), may counteract disease-associated molecular changes in BEC that otherwise prevent effective uptake of delivered Cu(btsc). Additionally, while other Cu compounds showed only modest transport through the AD iBEC monolayer, both Cu(ATSM) and Cu**L^1^** demonstrated increased permeability in AD iBEC indicating high potential for their translational application in AD patients. Simultaneously, control iBEC were more permissive to effective compound transport with the majority of tested metal compounds demonstrating increased permeability in monolayers formed by those cells.

It is important to note however that since no simple method can precisely distinguish the free metal (Cu or Ni) concentration and concentration of Cu/Ni bound to the (btsc) backbone, or freed (btsc) ligand itself within the cellular system *in vitro*, metal level measurement with ICP-MS was used here as an indirect measure of metal-(btsc)s transport. Therefore, it is not possible to unequivocally determine whether it was indeed an intact metal-(btsc) complex translocating through the monolayer of cells or only free metal that dissociated from the complex. Additionally, although various Cu(btsc) complexes were shown to be cell-permeable [69], performed ICP-MS analysis does not allow definition of whether cell-associated metal- (btsc) complex was indeed uptaken by the cells (reaching the cytosol) or remained only bound to the cell surface, therefore the transcellular transport of the compounds cannot be unequivocally confirmed. Despite this technical limitation, collectively those results provide clear evidence that modifications to metal–(btsc) structure modulate compound (or metal) transport at the brain barrier, with important disease-associated differences being observed between control and AD cells.

Although the exact molecular mechanisms responsible for the differential cellular accumulation and transport of distinct Cu(btsc) were not investigated in this study, a previous report identified opposite effects of Cu^II^(atsm) and Cu^II^(gtsm) on P-gp expression and function in the immortalised human brain endothelial cells *in vitro* [55] and demonstrated the regulatory effect of Cu^II^(atsm) on P-gp expression and function in murine BEC and mouse brain capillaries *in vivo* [97]. With P-gp being the primary drug efflux transporter at the BBB, it is possible that selected metal-(btsc) complexes exert a modulatory effect on P-gp expression and/or function and therefore escape P-gp-regulated efflux to reach higher intracellular concentrations in iBEC. Simultaneously, our previous analysis revealed differential expression and function of several drug transporters in the familial AD iBEC [40], [41], suggesting differences observed here in cell-associated accumulation of tested compounds can reflect disease-specific molecular alterations at the level of BBB transporters. However, both those hypotheses assume primarily the transcellular (as opposed to paracellular) transport of Cu compounds in this model which has not been experimentally validated.

Additionally, the differences observed in compound transport between control and AD cells in our model may suggest the role of presenilin-1 in Cu compound accumulation at the level of BBB. Since presenilins are known to promote cellular uptake of Cu [98], [99], *PSEN1-τ<E9* mutation present in our AD cells may drive aberrant Cu transport and therefore underlie decreased Cu compound permeability in AD iBEC. Interestingly, a similar observation was reported previously in the APP/PS1 mouse model, also harbouring mutant human presenilin 1, where structurally related Cu(btsc) compounds showed differential BBB permeability [68], suggesting the precise chemical design of small molecule drugs is crucial for their successful BBB transport in familial AD.

Although further mechanistic studies are required, our observations contribute to a further understanding of metal (btsc) effects at the BEC and demonstrate how small molecule compound physicochemical and structural characteristics correlate with its effective penetrability at the familial AD patient-specific BBB.

While Cu(btsc) complexes used as radiotracers were previously shown to successfully pass through the BBB, currently their brain uptake corresponds to ∼1 % of injected dose, which although correlating with sufficient uptake for brain imaging, is unlikely to achieve robust therapeutic effects in AD patients [53], [100]–[102]. It is therefore vital to continue efforts in designing and screening novel Cu(btsc) that may reach therapeutically relevant concentrations in the human brain to translate their neuroprotective effects from preclinical models to AD patients. Additionally, a recent study demonstrated that increased levels of brain Cu protect from cognitive decline in AD [103], further motivating the development of Cu- delivering agents such as Cu(btsc) as AD therapeutics. With hiPSC-derived platforms offering improved *in vitro* to *in vivo* correlation as compared to classically employed BBB models [31], compound delivery assays performed in our patient-specific model may prove to be highly predictive of Cu(btsc) transport in the patient brain and effectively inform metal drug candidate selection for future clinical trials.

### Neuroinflammation-like patient-derived BBB platform offers dual application in anti-inflammatory drug candidate screening

With numerous neuroinflammatory factors being involved in both the onset and the progression of AD [2], [3], [25], [30], our final goal was to evaluate the utility of our patient-derived platform to screen for the anti-inflammatory properties of novel metal compounds.

Importantly, although brain endothelial cells are not the primary immune cells in the brain, they are known to participate in inflammatory responses at the brain barriers both in health and disease [29], [104]–[106]. Respectively, a recent transcriptomic analysis revealed that many of the top AD-risk genes identified in genome-wide association studies (GWAS) which are microglia-specific in mice are in fact expressed at higher levels in human BEC and other vascular cell types, suggesting that in mice and humans, there is a partial transfer of risk genes and pathways associated with AD from microglia to the vasculature during the process of evolution [73]. This provides important evidence for the evolutionary-unique, human-specific role of BEC and cerebrovasculature in brain neuroimmunity in AD [73] and justifies efforts of anti-neuroinflammatory drug candidate screening in BEC. Simultaneously, disease-associated neuroinflammation has been shown to contribute to BBB dysfunction in AD [25], [30], [107]–[109], and a single nucleus transcriptomic analysis of AD patients’ brain revealed the upregulation of genes related to immune responses and cytokine secretion in BEC [72]–[74]; all suggesting that reducing inflammation in BEC could prove therapeutically useful in AD.

Interestingly, as opposed to BEC isolated from the AD patient’s brain [72]–[74], in our model we did not observe vast differences in the expression of inflammatory marker genes in AD iBEC as compared to control iBEC suggesting a lack of a strong disease-associated inflammatory phenotype in those AD cells *in vitro.* Intriguingly, recently reported transcriptomic analysis demonstrated striking molecular heterogeneity of human BEC found in different brain regions, distinct segments of the arteriovenous axis and disease stages, also in relation to immune responses [15], [73], [74]. Therefore it is possible our AD hiPSC-derived iBEC could represent the BEC subpopulation with only a moderately altered immune profile in AD, highlighting the complexity of BBB neuroinflammation in AD. Correspondingly, as it is not yet known which fraction of human *in vivo* BEC hiPSC-derived iBEC most closely represent, future comparative transcriptomic studies would be necessary to precisely map the position of iBEC within the human vasculature atlas and define their region-specific expression profile of genes related to cytokine production/immune response at various stages of AD.

Consequently, as AD iBEC did not present an intrinsic inflammatory profile in our model, we cultured those cells under proinflammatory conditions and validated the induction of their neuroinflammation-like phenotype *via* the panel of assays. Following treatment with TNFα and IFNγ, AD iBEC presented a characteristic increase in gene expression and cytokine secretion of several proinflammatory mediators, a decrease in barrier integrity and increased passive permeability to small molecule tracer, hence allowing us to test for anti-inflammatory properties of selected metal compounds in this model. As a result, our screen identified Cu**L^1^** as a strongly anti-neuroinflammatory compound which prevented the development of proinflammatory phenotype in AD iBEC in various assays tested. This presented also the first application of such a model for medium- to high-throughput screening of novel anti-neuroinflammatory drug candidates in familial AD iBEC.

Importantly, performing an anti-inflammatory drug candidate screen in such an established and validated platform offers dual benefits. Firstly, it presents a unique possibility to evaluate the anti-neuroinflammatory properties of novel compounds on the level of AD patient brain endothelial cells, previously not easily accessible to drug candidate screening efforts. With BBB neuroinflammation and neurodegeneration being closely linked in AD development and progression [25], [29], the discovery of such compounds may prove therapeutically useful in AD patients and lead to the identification of novel treatment avenues for AD. Correspondingly, Cu(btsc) were previously shown to act through multiple pathways involving anti-neuroinflammatory effects, the restoration of Cu homeostasis, inhibition of amyloid β and tau accumulation as well as the reduction in lipid peroxidation and ferroptosis [42], [43], [49], [56], with microglia, astrocytes and neurons being suggested as the major effector cells of their therapeutic action in the brain. Here, we detected potentially therapeutic immunomodulatory effects of selected Cu(btsc) in AD iBEC, implicating additional neuroprotective mechanisms that may be involved at the level of BBB. Intriguingly, Cu(btsc) were previously shown to exert therapeutic effects in preclinical models of AD, amyotrophic lateral sclerosis (ALS), and Parkinson’s disease (PD), as well as in first-in-human clinical trials targeting ALS and PD, all being neurodegenerative diseases with established roles for BBB neuroinflammation [110]–[114]. Correspondingly with BBB disruption becoming in itself an emerging drug target in neurodegeneration [115]–[118], further studies performed in our patient-derived model may support the ongoing efforts in identifying compounds that can improve cerebrovascular integrity in AD *via* active regulation of BEC, and accelerate the translation of those therapies to patients.

Secondly, our model allows to effectively test for drug candidate permeability at AD patient-derived BBB and pre-screen for its immunomodulatory effects within a single human-specific experimental set-up, rapidly identifying the most promising compounds for further assessment. Consequently, even for compounds whose therapeutic effects at the BBB are not of interest, our model can be effectively used for a first pre-evaluation of compound anti-inflammatory potential in human cells, supporting a more careful selection of BBB-permeable drug candidates to be then tested in e.g. microglia or astrocytes. With regards to that, implementing hiPSC-derived cells in anti-neuroinflammatory drug candidate screening platforms may be another important advantage as it allows for the differentiation of iBEC and classical immune cells such as microglia [119], [120] and astrocytes [57] from the same patient hiPSC. This may facilitate direct comparison of drug candidate responses in different cell types generated from hiPSC of the same patient, or lead to the development of multicellular isogenic models where iBEC can be co-cultured with other cells, such as induced-Astrocytes as previously demonstrated by us and others [22], [38], and compound BBB permeability and anti-inflammatory effects on brain parenchyma cells assessed within a single cell-culture well.

With this dual applicability, our iBEC model contributes to the advancement of metal-based therapies for AD via the effective identification of metal (btsc) with improved permeability and potential therapeutic anti-inflammatory activity in patient-derived cells. When validated against the library of novel compounds, experiments performed in this model identified Cu**L^1^** (Cu^II^(dtsm)) as a compound demonstrating high cellular uptake, permeability and anti-inflammatory effect in human AD BBB *in vitro*, which warrants its further testing in the context of AD-associated neuroinflammation in other brain cell models and *in vivo*.

### Limitations of the study

Overall our study demonstrates hiPSC-derived iBEC as an effective tool for modelling AD patient BBB *in vitro* and provides an alternative, validated platform for drug candidate permeability and efficacy screening. Although holding unprecedented potential, one limitation of disease modelling with hiPSC derived from independent human donors is their known inter-cell line variability [93], [121]. This allows for a more adequate representation of a heterogeneous patient population and therefore more translationally relevant preclinical drug candidate assessment (as opposed to i.e. immortalised cells lines or animal models). However, future studies utilising an increased number of patient cell-derived hiPSC lines may be required to confirm whether the observed here effects would be representative of a larger patient population. Similarly, with familial AD accounting for an estimated 5 % of all AD cases, expanding the presented here platform to our previously published sporadic AD iBEC model [38] may prove beneficial and increase model applicability to a wider AD patient cohort. It is also important to note that our study evaluated the cellular association and permeability of metal compounds at a single time point (2 h) which may not prove optimal for each compound given the known complexity of cellular [69] or brain [100], [101] accumulation profiles of structurally related Cu(btsc). Therefore, future studies incorporating multiple time points would be an important step towards understanding the temporal modes of action of Cu(btsc) at the AD BBB. Finally, to create a more physiologically relevant model, other BBB and parenchymal cells such as astrocytes or pericytes could be included. Incorporating elements of blood flow in the described here AD BBB model would also aid in achieving improved human BBB biomimicry *in vitro*.

With this potential for future assay-specific and disease subtype-specific modifications, our patient-derived model provides a versatile and flexible tool for routine BBB permeability testing offering unique advantages in the high-throughput drug candidate screening in AD.

## CONCLUSIONS

Early detection of BBB-permeable therapeutics may vastly accelerate successful drug development in AD. Our study exemplifies how hiPSC technology can be harnessed to assess BBB transport and anti-inflammatory effects of novel metal compounds in the familial AD context, offering a promising alternative to classically used preclinical BBB models.

Through practical validation of the established AD patient-derived iBEC model, our study identifies compound Cu^II^(dtsm) as a potential drug candidate with improved cell-associated accumulation and permeability in AD iBEC, and potentially therapeutic anti-neuroinflammatory activity at the AD BBB *in vitro*. Additionally, presented results suggest that Cu(btsc) complexes could be utilised as a new treatment approach to modulate neuroinflammation-associated BBB dysfunction in AD. Finally, by developing and testing a library of novel metal-(btsc) complexes we identify particular chemical structure modifications that facilitate low toxicity and improved compound transport at the AD BBB, supporting future design of small molecule therapeutics in AD.

Together, this disease- and patient-relevant model may serve as an innovative drug candidate screening platform with higher translational significance and improved *in vivo* predictivity as compared to traditionally employed BBB permeability assays. When applied together with other pharmacokinetic and pharmacodynamics methods, it can aid in the early identification of CNS-active, -permeable and non-cytotoxic compounds, significantly contributing to the therapeutic success of drugs targeting AD.

## MATERIALS AND METHODS CELL MODELS

### Human Umbilical Vein Endothelial Cells (HUVEC) culture and immunofluorescence characterisation

Primary human umbilical vein endothelial cells (HUVEC) (Life Technologies) were cultured in 75 cm^2^ flasks in Endothelial Cell Growth Media (Sigma) under normoxia conditions (37 °C, 5 % CO_2_).

For immunofluorescence (IF) characterisation HUVEC were cultured on coverslips coated with 10 μg/ml human fibronectin until reaching 100 % confluency and forming a cobblestone-like monolayer. Cells were then fixed with 4 % paraformaldehyde (PFA; Sigma) for 15 min at room temperature (RT) and washed with phosphate-buffered saline (PBS) and IF was performed as follows: cells were permeabilised for 10 min with 0.3 % Triton-X (Sigma) and then blocked for 1 h at RT with 2 % bovine serum albumin (BSA)/2 % normal goat serum (GS) in PBS. The primary antibody for vascular endothelial (VE)-cadherin (**Table S1**) was diluted at 1:100 in a blocking solution and incubated overnight at 4 °C. After 24 h, cells were washed with PBS and secondary antibodies (Alexa Fluor-488; **Table S1**) diluted in blocking solution (1:250) were incubated on the cells for 1 h at RT in the dark. Cells were then washed with PBS, Hoechst (1:5000 in PBS) counterstain was performed to visualise cell nuclei and cells mounted with ProLong Gold Antifade (ThermoFisher Scientific). Images were obtained at 20X magnification using a Zeiss AxioScop2 microscope.

### Human induced pluripotent stem cells (hiPSC) culture and immunofluorescence characterisation

Previously published and characterised human induced pluripotent stem cell (hiPSC) were obtained from the University of Eastern Finland [57] and the University of Melbourne [40]. hiPSC lines: 1 x healthy control line (referred to as HDFa), 2 x *PSEN1-τ<E9* mutant AD line, 2 x isogenic control to *PSEN1-τ<E9* were used in this study (**Table 1**). All hiPSC were expanded on human recombinant vitronectin in StemFlex^TM^ media (ThermoFisher Scientific) under hypoxia conditions (37 °C, 5 % CO_2_, 3 % O_2_). During initial expansion, hiPSC were passaged with 0.5 mM ethylenediaminetetraacetic acid (EDTA, Life Technologies) in PBS and cryopreserved in 10 % dimethyl sulfoxide (DMSO, Sigma) in StemFlex^TM^ media. Karyotype analysis was performed for HDFa, 1 x *PSEN1 τ<E9* mutant AD line and 1 x isogenic control to *PSEN1-τ<E9.* All hiPSC lines tested showed a normal karyotype, containing 22 pairs of autosomal chromosomes and one pair of sex chromosomes (46, XX) (data not shown).

For IF characterisation, cells were fixed with 4 % PFA for 15 min, rinsed with PBS and permeabilised with 0.3 % Triton-X for 10 min. Cells were then blocked for 1 h at RT with 2 % BSA/2 % GS in PBS. Primary antibodies for Nanog and SOX2 (**Table S1**) were diluted at 1:100 in blocking solution and incubated overnight at 4 °C. The next day, PBS washes were performed and secondary antibodies (Alexa Fluor-488, or Alexa Fluor-647; **Table S1**) were diluted in blocking solution (1:250) and incubated on the cells for 1h at RT in the dark. Cells were then washed with PBS and Hoechst (1:5000) counterstain was performed. Coverslips with cells were mounted with ProLong Gold Antifade. Images were obtained at 20X magnification using a Zeiss 780 confocal microscope.

### Induced brain endothelial-like cell (iBEC) differentiation

To establish a patient-derived BBB model, hiPSC were differentiated towards brain endothelial-like cell phenotype following previously published protocols [38], [40], [58]. At all stages of iBEC differentiation cells were cultured under normoxia conditions (37 °C, 5 % CO_2_). To initiate iBEC differentiation, hiPSC were detached and singularised with Accutase (Life Technologies) and plated on human embryonic stem cells (hESC)-qualified Matrigel (Corning) coated 6-well culture plates in StemFlex^TM^ media supplemented with 10 μM Rho-associated kinase inhibitor (iROCK) at previously optimised [40] plating density of 2.0 x 10^4^ cell/well (HDFa line) or 2.5 x 10^4^ cell/well (isogenic corrected and AD lines). After 3 days, culture medium was changed to unconditioned media (UM) consisting of DMEM/F12+GlutaMAX (Life Technologies), 20 % KnockOUT serum replacement (Life Technologies), 1 x non-essential amino acids (Life Technologies) and 0.1 mM β-mercaptoehtanol (Sigma) to induce neural and endothelial progenitor co-differentiation. Following 6 days in UM, culture media was replaced with endothelial cell media (EC; Life Technologies) supplemented with 2 % B27 (Life Technologies), 20 ng/ml basic fibroblast growth factor (FGFb; Peprotech) and 10 μM retinoic acid (RA). Cells were maintained in supplemented EC+B27 for 2 days, after which cells were detached and singularised with Accutase, and replated on collagen IV from human placenta (Sigma) and human plasma fibronectin (Life Technologies) coated plastic culture plates or Ø 0.4 μm pore polyester Transwell inserts (Corning) at plating density specific for culture vessel (**Table S3**). The day after subculturing cells to collagen IV/fibronectin plates, cell media was changed to EC+B27 (without FGFb and RA) and cultured for one more day. All experiments described in this study were performed 48 h following subculturing on collagen IV/fibronectin.

### iBEC immunofluorescence characterisation

For IF characterisation, iBEC were grown on plastic coverslips coated with 80 μg/ml collagen IV and 20 μg/ml fibronectin. 48 h after subculturing, cells were rinsed with PBS and fixed with ice-cold 100 % methanol (MeOH) for 5 min at -20 °C or 4 % PFA for 15 min at RT. Next cells were permeabilised with 0.3 % Triton-X for 10 min and blocked for 1 h at RT with 2 % BSA/2 % GS in PBS. Primary antibodies for occludin, claudin-5, ZO-1 and Glut-1 (**Table S1**) were diluted 1:100 in a blocking solution and incubated overnight at 4°C. The next day, PBS washes were performed and secondary antibodies (Alexa Fluor-488 or Alexa Fluor-647; **Table S1**) were diluted 1:250 in a blocking solution and incubated on the cells for 1 h at RT in the dark. Afterwards, cells were washed with PBS and Hoechst (1:5000) counterstain was performed. The coverslips with cells were mounted with ProLong Gold Antifade. Images were obtained at 20X magnification using a Zeiss 780 confocal microscope.

### Transendothelial electrical resistance (TEER) measurement

Barrier integrity of generated iBEC was characterised by measuring transendothelial electrical resistance (TEER) across iBEC monolayer using the EVOM2 or EVOM3 Volt/Ohmmeter (World Precision Instruments) in 24-well, 6.5 mm Transwell with 0.4 μm pore polyester membrane insert (Corning). Before the measurement, TEER electrodes were sterilised and immersed in warm EC+B27 media for temperature equilibration. TEER was then measured in 3 areas per Transwell and averaged. Resistance of the blank (no-cells) Transwell was subtracted and then multiplied by the surface area of the Transwell membrane (0.33 cm^2^) for calculation of the final TEER values (Ohm x cm^2^).

## METAL BIS(THIOSEMICARBAZONE) COMPOUNDS SYNTHESIS

### General

^1^H and ^13^C{^1^H} spectra were recorded using a Varian FT-NMR 400 spectrometer (Varian). All ^1^H NMR spectra were acquired at 400 MHz and ^13^C{^1^H} spectra were acquired at 101 MHz. The reported peaks were all referenced to solvent peaks in the order of parts per million at 25 °C. Microanalysis measurements were carried out by The Campbell Microanalytical Laboratory in the Department of Chemistry, University of Otago, Union Place, Dunedin, New Zealand. Analytical HPLC were performed on Agilent 1200 series HPLC system fitted with an Alltech Hypersil BDS – C18 column (4.6 × 150 nm, 5 µm). The mobile phase was a gradient consisting of Solvent A (0.1 % TFA in H_2_O) and Solvent B (0.1 % TFA in CH_3_CN) from 0 to 100 % B over 25 min and UV detection at λ 220, 254, 275 and 350 nm. ESI-QTOF MS was collected on an Exactive Plus Orbitrap Infusion mass spectrometer (Exactive Series, 2.8 Build 268801, ThermoFisher Scientific). Analysis was performed using Xcalibur 4.0.27.10 (ThermoFisher Scientific).

### Chemical Synthesis

Cu(ATSM), Ni(ATSM), Cu(DTSM) (CuL^1^), Cu(DTSE) (CuL^2^) [95], [122] and H_2_ATSM/M_2_ [123], [124] were synthesized as reported previously. H_2_DTSM/M_2_ and H_2_DTSE/M2 were prepared by modification of previously reported procedure [123] where dipropionyl-mono-4-methyl-3- thiosemicarbazone or dipropionyl-mono-4-ethyl-3-thiosemicarbazone [68] were reacted with one equivalent of 4,4-dimethyl-3-thiosemicarbazide in dimethyl formamide in the presence of acetic acid.

Ligands H_2_L^3-10^ were prepared by modification of a reported procedure [123] where either ATSM/M_2_, H_2_DTSM/M_2_ or H_2_DTSE/M_2_ are reacted with the requisite primary amine (see **Scheme 1**). All the ligands were isolated in yields of 30 – 80 % (depending on the solubility of the ligand in acetonitrile), and were characterised by ^1^H and ^13^C{^1^H} spectroscopy and electrospray ionisation mass spectrometry. The metal complexes were prepared by the reaction of the ligand, H_2_L*^x^*, with one equivalent of copper acetate monohydrate to give CuL^1-10^ (or nickel acetate monohydrate to give NiL^3^) and heating the mixtures in ethanol at reflux for 4 hours (**Scheme 1**). For the synthesis of CuL^1-7^: Allowing the reaction mixture to cool to room temperature resulted in precipitation of brown-red solids that were collected by filtration, washed with cold ethanol and diethyl ether to allow isolation of CuL^1-7^ in ∼70 % yield. For the synthesis of CuL^8-10^: The reaction mixture was evaporated to dryness under reduced pressure, the solid was then dissolved in dichloromethane and addition of *n*-pentane resulted in the precipitation of dark red solids that were collected by filtration, washed with *n*-pentane and dried in vacuo to allow isolation of CuL^8-10^ in ∼ 60 % yield. All the copper(II) complexes were characterised by electrospray ionisation mass spectrometry, reversed phase HPLC (>98% purity) and microanalysis.

**Scheme 1.**
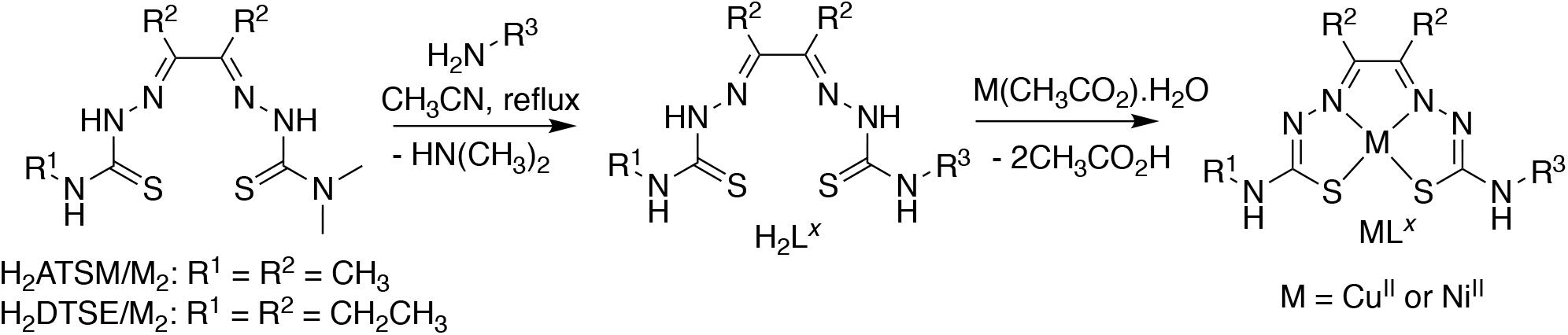
General synthetic scheme for the synthesis of ligands, H_2_L^1^-H_2_L^10^, and metal complexes ML^1-10^ (where M = Cu^II^ or Ni^II^).

A representative procedure for the synthesis of H_2_L^4^ and CuL^4^ is given below:

### Synthesis of H_2_L^4^

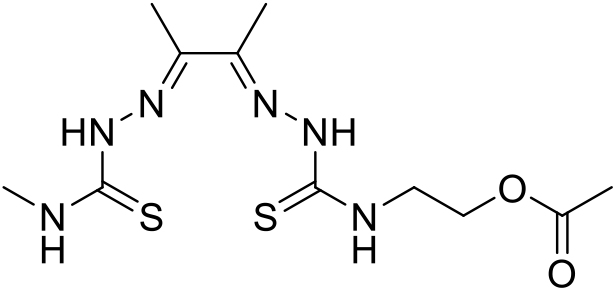

To a suspension of H_2_ATSM/M_2_ (0.5 g, 1.8 mmol) in acetonitrile (50 mL) was added aminoethyl acetate (0.23 g, 2.2 mmol). The mixture was heated at reflux for 24 h and then allowed to cool to ambient temperature. The precipitate that formed was collected by filtration, washed with acetonitrile, water and diethyl ether to give product as a pale pink solid (0.51 g, 1.5 mmol, 83 %). ^1^H NMR (400 MHz, DMSO-d_6_, δ) 10.35 (s, 1H), 10.23 (s, 1H), 8.46 (t, J = 5.7 Hz, 1H), 8.38 (d, J = 4.5 Hz, 1H), 4.20 (t, J = 5.7 Hz, 2H), 3.81 (q, J = 5.7 Hz, 2H), 3.02 (d, J = 4.5 Hz, 4H), 2.21 (d, J = 3.3 Hz, 6H), 2.01 (s, 3H). ^13^C NMR (101 MHz, DMSO-d_6_, δ) 178.48, 178.28, 170.43, 148.57, 147.78, 62.05, 42.65, 31.22, 20.79, 11.81, 11.65. MS(ESI/O-TOF) (m/z): Cald for [C_11_H_20_N_6_O_2_S_2_+H]^+^, 333.1162; found, 333.1163. HPLC R_t_ = 8.4 min. Anal Calcd C, 39.74; H, 6.06; N, 25.28. Found C, 39.12; H, 5.77; N, 24.91

### Synthesis of CuL^4^

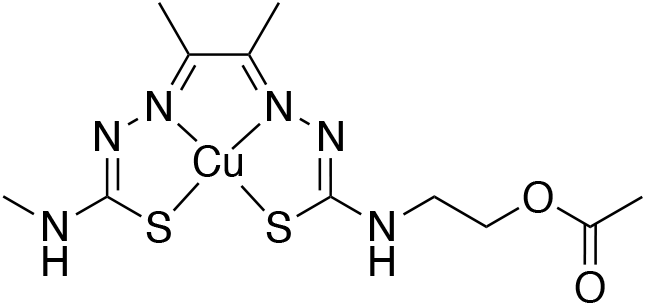

To a suspension of H_2_L^4^ (0.2, 0.6 mmol) in ethanol (10 mL) was added copper acetate mono hydrate (0.12, 0.60 mmol). The mixture was heated at reflux for 20 h and was then allowed to cool to ambient temperature. The precipitate that formed was collected by filtration, washed with ethanol, water and diethyl ether to give CuL^4^ as a brown powder (0.17 g, 0.43 mmol, 72%). MS(ESI/O-TOF) (m/z): Cald for [C_11_H_18_CuN_6_O_2_S_2_+H]^+^, 394.0301; found, 394.0301. HPLC: R_t_ = 6.1 min. Anal. Calcd for C_11_H_18_CuN_6_O_2_S_2_: C, 33.54; H, 4.61; N, 21.33. Found: C, 33.11; H, 4.67; N, 21.08.

## CELL STIMULATION AND/OR TREATMENT

### MTT cytotoxicity assay

Cu and Ni based compounds were diluted in 100 % DMSO (Sigma). Cells in a 96-well plate (HUVEC: 3 × 10^3^ cells/well; iBEC: 1 × 10^5^ cells/well) were treated with increasing concentrations of Cu- and Ni-based compounds (HUVEC: 0, 0.1, 0.5, 1.0, 1.5, 2.0, 2.5, and 3.0 μM; iBEC: 0, 0.5 and 1.0 μM) for 24 h. (3-(4,5-dimethylthiazol-2-yl)-2,5- diphenyltetrazolium bromide (MTT, Sigma) reagent was added to each well, and the cells were incubated in humidified incubator for 4 h at 37 °C. Subsequently, solubilisation solution (10 % Triton-X 100 in acidic isopropanol (0.1 M HCl)) was added to wells and incubated overnight at RT on orbital shaker. The next day, absorbance was measured at 570 nm with a microplate reader (Biotek Synergy H4) and % of viable cells was calculated as compared to the untreated control. Vehicle (DMSO)-only controls were included for the two highest drug concentrations tested (corresponding to DMSO content equivalent to 2.5 and 3.0 µM of a drug treatment for HUVEC and 0.5 and 1.0 µM of a drug treatment for iBEC, respectively).

### Metal cell-associated accumulation and permeability analysis

To analyse Cu and Ni cell-associated accumulation and permeability, iBEC were cultured on Transwell inserts with polyester membranes containing 0 0.4 μm pores. 48 h after subculture TEER was measured and only cells exhibiting adequate TEER indicating complete monolayer formation were included in the following experiment. Following TEER assessment, cells were allowed to recover for 1 h in the incubator. Tested compounds were then added to the top chamber of Transwells at 0.5 μM and incubated for 2 h. Both untreated and vehicle-only treated controls were included. Next, cell pellets and cell culture media from the top and bottom chambers of a Transwell system were collected for inductively coupled plasma mass spectrometry (ICP-MS) analysis of metal levels, performed at Biometals Facility at Florey Institute of Neuroscience and Mental Health. For media sample analysis, media collected from a single Transwell was considered an independent replicate. Due to the low number of cells grown on a single Transwell membrane, for cell pellet analysis, cells grown on three Transwells were combined into one tube prior to the pelleting and considered as an independent replicate. For analysis of metal levels in media sample, a 900 μl of diluent (1 % nitric acid) was added to each media sample (100 μl), to equal 1 ml of final volume. For the analysis of metal levels in collected cell pellet samples, cell pellets were lyophilized, 30 μl of concentrated 65 % nitric acid (Suprapur, Merck) was added and allowed them to digest overnight at RT. The samples were then heated at 90 °C for 20 min using a heating block to complete the digestion. The reduced volume after digestion was ∼20 μl. Next, 580 μl of 1 % (v/v) of nitric acid diluent was added to equal 0.6 ml of final volume. Measurements were made using an Agilent 7700 series ICP-MS instrument under routine multi-element operating conditions using a Helium Reaction Gas Cell. The instrument was calibrated using 0, 5, 10, 50, 100 and 500 parts per billion (ppb) of certified multi-element ICP-MS standard calibration solutions (ICP-MS-CAL2-1, ICP-MS-CAL-3 and ICP-MS-CAL-4, Accustandard) for a range of elements. A certified internal standard solution containing 200 ppb of Yttrium (Y89) was used as an internal control (ICP-MS-IS-MIX1-1, Accustandard). The concentration of Cu and Ni samples in cell pellet and media samples were calculated as follows: cell pellets: [μmol/L] = (raw ppb value x final sample volume/molecular weight of the element) media samples: [μmol/L] = (raw ppb value x dilution factor/molecular weight of the element). Cu and Ni levels in media samples were expressed as concentration [μmol/L] and cell pellet Cu and Ni levels were normalised to concentration of Mg per each sample for the uptake analysis. Mg was used as an internal divalent control metal to standardise Cu and Ni values, and the levels of Mg between analysed samples were not statistically different.

### TNFα / IFNγ treatment and metal compound treatment

To study iBEC response to inflammatory stimuli, iBEC were treated with tumour necrosis factor alfa (TNFα, ThermoFisher Scientific; 20 ng/ml) with interferon gamma (IFNγ, ThermoFisher Scientific; 30 ng/ml) for 24 h. For co-treatment experiments, metal compounds were diluted in cell culture media and added to cells at 0.5 μM for 24 h with or without 20 ng/ml TNFα/ 30 ng/ml IFNγ. Respective vehicle-only controls (DMSO for metal compound treatment and 40 mM Tris buffer (Sigma-Aldrich) for TNFα and IFNγ treatment) were included.

### Lactate dehydrogenase (LDH) cytotoxicity assay

LDH cytotoxicity assay was performed to assess TNFα/IFNγ effect on cell viability. iBEC were cultured in 48 well plates and exposed to TNFα/IFNγ for 24 h and the levels of LDH enzyme in the collected media were determined using CyQUANT LDH Cytotoxicity Assay (ThermoFisher Scientific) following manufacturer instructions. To determine maximum lactate dehydrogenase (LDH) release (LDH_Max_) cells from selected wells were treated with the Lysis Buffer (10 % TritonX in PBS) for 30 min. The absorbance was measured at 490 nm and 680 nm using a plate reader (Biotek Synergy H4). To determine LDH activity, the 680 nm absorbance value (background) was subtracted from the 490 nm absorbance and the % of LDH_max_ was calculated in the analysed samples. Data are presented as fold change in relative LDH release corresponding to vehicle-treated control.

### IL6 and MCP1 enzyme-linked immunosorbent assay (ELISA)

To detect interleukin-6 (IL6) and monocyte chemoattractant protein-1 (MCP1, CCL2) cytokine secretion by iBEC following TNFα/IFNγ and drug stimulation, iBEC were cultured in 48 well plates, treated with TNFα/IFNγ with or without the tested drug for 24 h and cell culture media collected for ELISA. Respective vehicle-only controls were included. Immediately post- collection, samples were centrifuged at 300 x *g* for 3 min to remove cell debris and supernatant stored at -80 °C prior to analysis. Human IL-6 uncoated ELISA (Invitrogen) and human CCL2 (MCP1) uncoated ELISA (Invitrogen) were used as per manufacturer instruction. All incubation steps were performed using a microplate shaker.

In brief, 96 well Coat Corning™ Costar™ 9018 ELISA plates were coated with capture antibody (pre-titrated, purified anti-human IL6 and purified anti-human CCL2 antibody for IL6 and MCP1 ELISA respectively) in coating buffer overnight at 4 °C. Plates were then washed with Wash Buffer (0.05 % Tween®20 (Sigma-Aldrich) in PBS) and blocked with ELISA/ELISPOT Diluent for 1 h at RT. Next, human IL6 and CCL2 standards were freshly prepared, plates were washed with Wash Buffer and samples and Standards serial dilutions added to appropriate wells. ELISA/ELISPOT Diluent was added to the blank well and plates incubated for 2 h in RT. Plates were then washed with Wash Buffer and Detection Antibody (pre-titrated, biotin-conjugated anti human IL6 antibody and biotin-conjugated anti-human CCL2 antibody for IL6 and MCP1 ELISA respectively) added to the wells. Following 1 h incubation, plates were washed with Wash Buffer and incubated with Streptavidin-horseradish peroxidase (HRP) enzyme for 30 min. Plates were subsequently washed with Wash Buffer and incubated with tetramethylbenzidine (TMB) substrate solution for 15 min at RT. Reaction was then stopped by adding 2 N H_2_SO_4_ Stop Solution (Sigma) and absorbance was measured at 450 nm and 570 nm using a plate reader (Biotek Synergy H4). Obtained IL6 and CCL2 standard curves were used to determine the concentration of IL6 and MCP1 in analysed samples.

### iBEC barrier properties assessment following TNFα / IFNγ treatment and metal compound treatment

To analyse effects of TNFα / IFNγ and metal compound treatment on iBEC barrier properties, iBEC were cultured on Transwell inserts with polyester membranes containing 0 0.4 μm pores. 48 h after subculture TEER was measured and only cells exhibiting adequate TEER indicating complete monolayer formation were included in the following experiment.

To assess the integrity of iBEC monolayer, cells were treated with vehicle or TNFα/IFNγ and Cu**L^1^** (0.5 μM) and Cu(ATSM) (0.5 μM) alone or in combination, and TEER was measured at 24 h after the start of the treatment as described above. Fold changes in TEER were calculated as compared to vehicle-treated control.

To assess the passive permeability of iBEC monolayer following TNFα/IFNγ and Cu**L^1^** (0.5 μM) and Cu(ATSM) (0.5 μM) treatments, fluorescein isothiocyanate (FITC)-conjugated dextran molecule of 3–5 kDa (Sigma) was added at 0.5 mg/ml to the top chamber of the Transwell insert at 24 h after the start of the treatment. Following 1 h incubation with 5 kDa dextran, the top and bottom chamber media were collected for spectrofluorometric analysis at 490 nm excitation/520 nm emission using a fluorescent plate reader (Biotek Synergy H4). Clearance volume describing dextran permeability was calculated as described previously [38], [58] following formula:

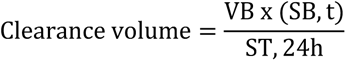

where VB is the volume of the bottom chamber (800 μl); SB,t is the corrected signal of the bottom chamber at the time, t; ST,24h is the signal of the top chamber at 24 h. Permeability results were presented as fold changes in clearance volume and compared to vehicle-treated control.

## GENE EXPRESSION STUDIES

### RNA extraction

For cell phenotype characterisation, hiPSC and iBEC were cultured under normal conditions prior to RNA collection. For the study of the inflammatory response, iBEC were cultured on 48-well plates and exposed to TNFα/IFNγ alone or in combination with metal compound (0.5 µM) treatments for 24 h prior to RNA collection. Respective vehicle-only controls were included.

For RNA collection, cells were rinsed with PBS, lysed with TRIzol^TM^ reagent (ThermoFisher Scientific) and were stored in -80 °C prior to RNA extraction. Total RNA was extracted using the Direct-zol RNA Miniprep Kit (Zymo Research) according to the manufacturer’s instructions and treated in-column with DNase I. In brief, sample lysed in TRIzol was thawed on ice and equal volume of analytical grade 100 % ethanol (EtOH, Chem-Supply) was added and mixed. The mixture was then transferred on Zymo-Spin^TM^ IICR Column and centrifuged at 10,000 x *g* for 1 min. The column was washed with RNA Wash Buffer and sample incubated with 0.375 U/μl of DNAse I for 15 min. Next, the column was washed two times with Direct-zol^TM^ RNA PreWash Buffer and RNA Wash Buffer before final elution in DNAse/RNAse free water. Isolated RNA quality and quantity was measured using NanoDrop^TM^ Spectrophotometer.

### cDNA synthesis and quantitative PCR (qPCR)

For quantitative polymerase chain reaction (qPCR) studies, 50 ng (for assays where cells cultured in 48-well plates) or 150 ng (for cells cultured in 24-well plates) of total RNA was converted to complementary DNA (cDNA) using SensiFAST^TM^ cDNA synthesis kit following manufacturer instructions (Bioline). For cDNA synthesis reaction mix containing adequate volumes of RNA, 2 μl of 5 x TransAmp Buffer, 0.5 μl of Reverse Transcriptase (RT) enzyme and DNAse/RNAse free water was prepared. Appropriate no-RNA template and no-RT control reactions were included. 384 well plate containing reaction mixture was centrifuged at 300 x *g* for 1 min and cDNA synthesis performed in a thermal cycler (T100, Bio-Rad Laboratories) using following program: 25 °C for 10 min (primer annealing), 42 °C for 15 min (reverse transcription), 85 °C for 5 min (inactivation), 4 °C hold.

Subsequently, cDNA was diluted 1:10 in DNAse/RNAse free water to generate working solution and qPCR performed using SensiFAST™ SYBR® Lo-ROX Kit following manufacturer instructions (Bioline). For qPCR, a reaction mix of 2 μl cDNA template, 2.2 μl H_2_O, 400 nM of gene-specific primers (**Table S2**) and 5 μl of SensiFAST^TM^ SYBR® Lo-ROX reagent was prepared. The qPCR run was performed as triplicate for each sample on QuantStudio^TM^ 5 Real-Time PCR system with following run conditions: 2 min at 95 °C followed by 40 cycles of 5 s at 95°C and 30 s at 60°C. Ct values were normalised to Ct values of *18S* endogenous control (ΔCt values). *18S* housekeeping gene expression was found to be consistent across cell lines, conditions and time points. Standard deviation (SD) was calculated for each technical triplicate and samples with SD > 0.5 excluded from analysis. ΔΔCt values were calculated as 2(^-ΔCt^) and presented as ΔΔCt multiplied by 10^6^ or fold changes in ΔΔCt x 10^6^. Technical replicates were averaged per sample for statistical analysis.

## STATISTICAL ANALYSIS

Statistical analysis was performed using GraphPad Prism version 9.4.0. Data were analysed using an unpaired t-test with Welch’s correction when a comparison between two groups was being investigated or one-way ANOVA followed by post-hoc tests when comparisons between three or more groups were analysed, with *p* value of less than < 0.05 considered statistically significant. For data identifying as potential outliers, Z-score was calculated for each value and values with Z-score exceeding 2 or -2 (indicating 2 standard deviations (SD) above or below the mean) were identified as outliers and excluded from analysis. Results are shown as mean ± SEM unless specified differently in the figure legends. The number of biological (N, hiPSC or iBEC lines) and independent (n) replicates used for each experiment are specified in figure legends.

## SUPPLEMENTARY MATERIALS

Supplementary figures:

**Figure. S1.** Effects of DMSO-only (vehicle) on the viability of the human umbilical vein endothelial cells (HUVEC) and control, and AD, induced brain endothelial-like cells (iBEC).

**Figure. S2.** Characterisation of the healthy donor, isogenic-corrected control and *PSEN1- ΔE9* familial AD hiPSC lines.

**Figure. S3.** Schematic flow of hiPSC-derived iBEC differentiation.

**Figure. S4.** Characterisation of the healthy donor, isogenic-corrected control and *PSEN1-ΔE9* familial AD iBEC.

**Figure. S5.** Comparison of the cytotoxic effects of selected metal compounds between control and AD iBEC.

**Figure. S6.** Baseline expression of proinflammatory, oxidative stress and endothelial cell junctional marker genes in control and AD iBEC.

**Figure. S7.** Effects of TNFα/IFNγ- stimulation on control iBEC phenotype.

**Figure. S8.** Comparison of AD iBEC responses to CuL^1^, Cu(ATSM) and Ni(ATSM) treatments.

### Supplementary tables

**Table S1.** Antibodies used in the study.

**Table S2.** Primer sequences used in the study.

**Table S3.** Coating solution concentration and cell plating density defined per specific culture plate type utilised during iBEC purification step.

**ABBREVIATIONS:** AD: Alzheimer’s disease; ALS: amyotrophic lateral sclerosis; BBB: blood-brain barrier; BEC: brain endothelial cell; (btsc): bis(thiosemicarbazone); *CDH5*: VE-cadherin; *CLDN5*: claudin-5; CNS: central nervous system; Cu: copper; ELISA: enzyme-linked immunosorbent assay; hiPSC: human induced pluripotent stem cell; human umbilical vein endothelial cells (HUVEC); iBEC: induced brain endothelial-like cell; ICP-MS: inductively coupled plasma mass spectrometry; IL-interleukin; IFNγ: interferon γ; L: ligand; LDH: lactate dehydrogenase; MCP1: monocyte chemoattractant protein-1; MTT: 3-(4, 5-dimethyl thiazol- 2)-2, 5-diphenyltetrazolium bromide; *OCLN*: occludin; PAMPA: parallel artificial membrane permeability assay; PD: Parkinson’s disease; P-gp: P-glycoprotein; *PSEN1:* presenilin-1; qPCR: quantitative polymerase chain reaction; TEER: trans-endothelial electrical resistance; *TJP1* – zonula occludens 1; TNFα: tumour necrosis factor α; veh. ctrl: vehicle control;

## Acknowledgements

We acknowledge the QIMR Berghofer Medical Research Institute Microscopy Facility team for their assistance and the Florey Institute Biometals Facility for sample processing. We thank Dr Carolin Offenhauser for the provision of HUVEC, prof. Jose M. Polo for the provision of HDFa hiPSC line and Dr Romal Stewart for the critical reading of the manuscript. Graphical elements of figures were created with Biorender.com.

## CRediT authorship contribution statement

**Joanna M. Wasielewska**: conceptualisation, methodology, investigation, formal analysis, visualisation, writing - original draft, writing - review & editing; **Kathryn Szostak**: methodology, resources; **Lachlan E. McInnes**: methodology, resources; **Hazel Quek**: methodology, resources, formal analysis, writing - review & editing; **Juliana C. S. Chaves**: methodology; **Jeffrey R. Liddell**: methodology; **Jari Koistinaho**: resources (provision of hiPSC lines); **Lotta E. Oikari**: conceptualisation, methodology, supervision; **Paul S. Donnelly**: conceptualisation, methodology, resources, supervision, writing - review & editing; **Anthony R. White**: conceptualisation, writing - review & editing, supervision, project administration, funding acquisition. All authors reviewed and approved final version of the manuscript.

