## Supplementary Materials for "A patient-derived blood-brain barrier model for screening copper bis(thiosemicarbazone) complexes as potential therapeutics in Alzheimer’s disease"

Wasielewska *et al*.

### SUPPLEMENTARY FIGURES

Wasielewska *et al*.

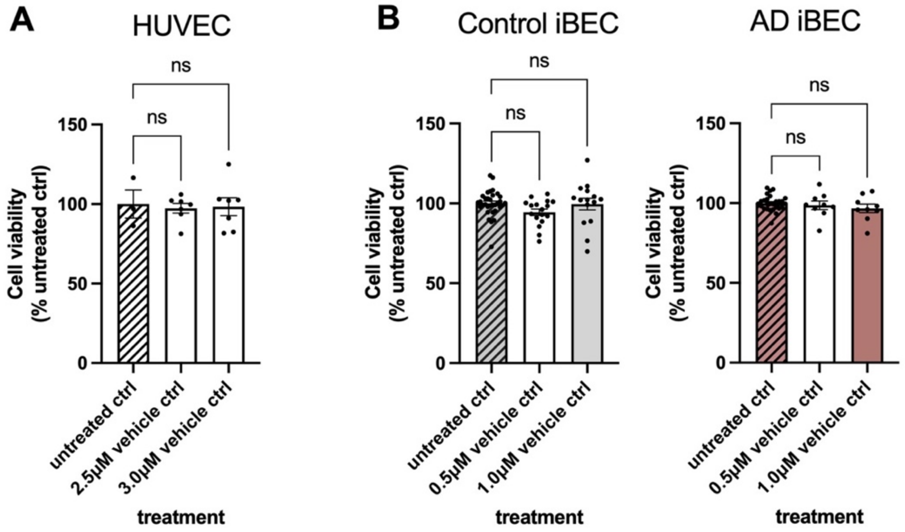

**Figure S1. Effects of DMSO-only (vehicle) on the viability of the human umbilical vein endothelial cells (HUVEC) and control, and AD, induced brain endothelial-like cells (iBEC).** (**A**) HUVEC and (**B**) iBEC were treated with DMSO-only (vehicle) at concentrations corresponding to the two highest concentrations of compound tested in Figure 1C and Figure 2D-D' (DMSO content corresponding to compound treatment at 2.5 μM and 3.0 μM for HUVEC, and 0.5 μM and 1.0 μM for iBEC was tested, respectively) and cell viability assessed with MTT assay. Cell viability is shown as % of viable cells compared to untreated control (ctrl). In (A) a minimum of n=3 independent replicates per condition; in (B) Control iBEC: N=3 lines, AD: N=2 lines; a minimum of n=3 independent replicates per line. Data are presented as mean ± SEM. Statistical analysis in (A-B) was performed using one-way ANOVA with Dunnett's test.

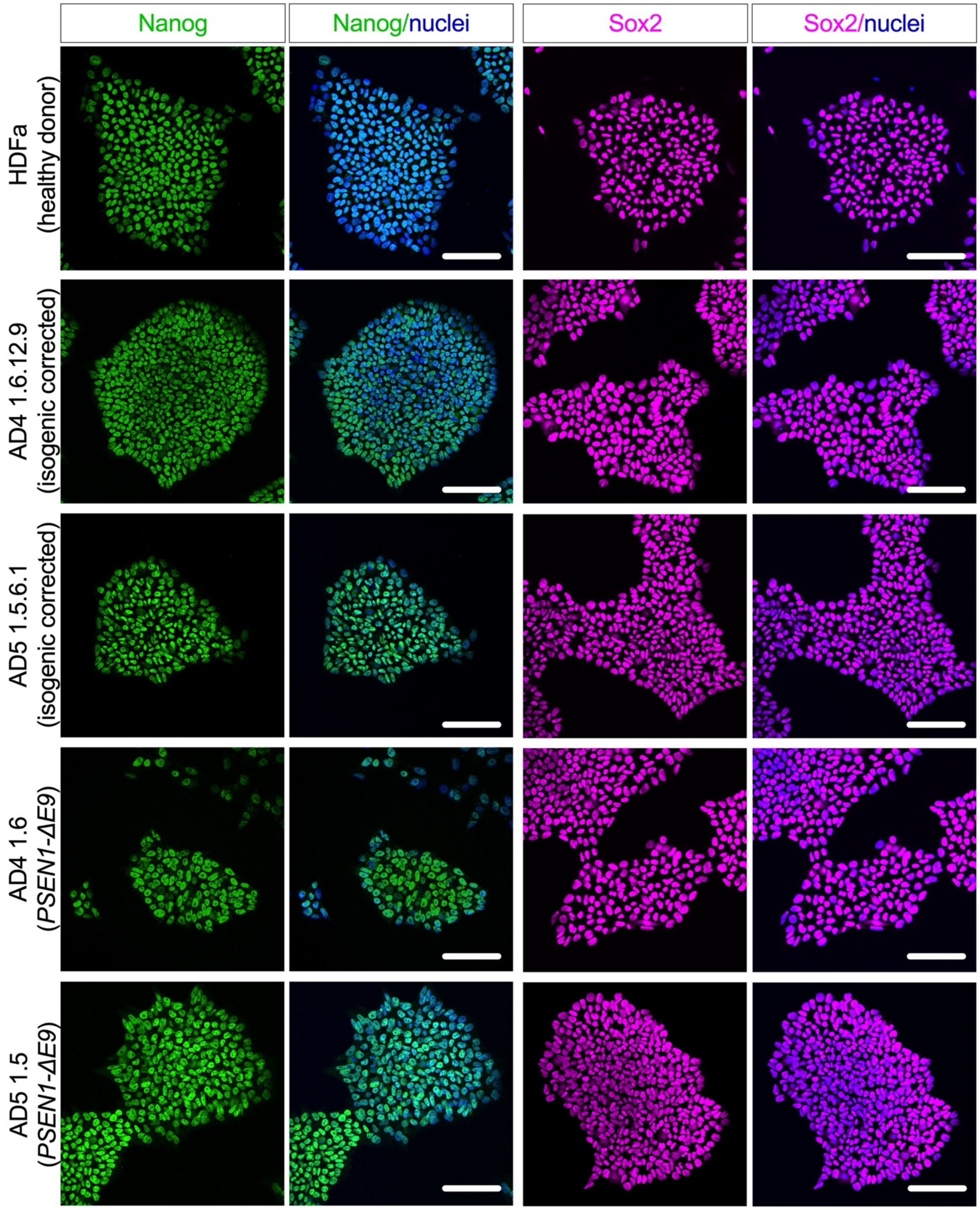

**Figure S2. Characterisation of the healthy donor, isogenic-corrected control and *PSEN1-ΔE9* familial AD hiPSC lines.** Representative immunofluorescence images of the pluripotency markers Nanog (green) and Sox2 (magenta) with Hoechst (blue) nuclear counterstaining in HDFa, isogenic-corrected control AD4 1.6.12.9 and AD5 1.5.6.1, and *PSEN1-ΔE9* AD4 1.6 and AD5 1.5 hiPSC lines. Scale bar, 100 µm.

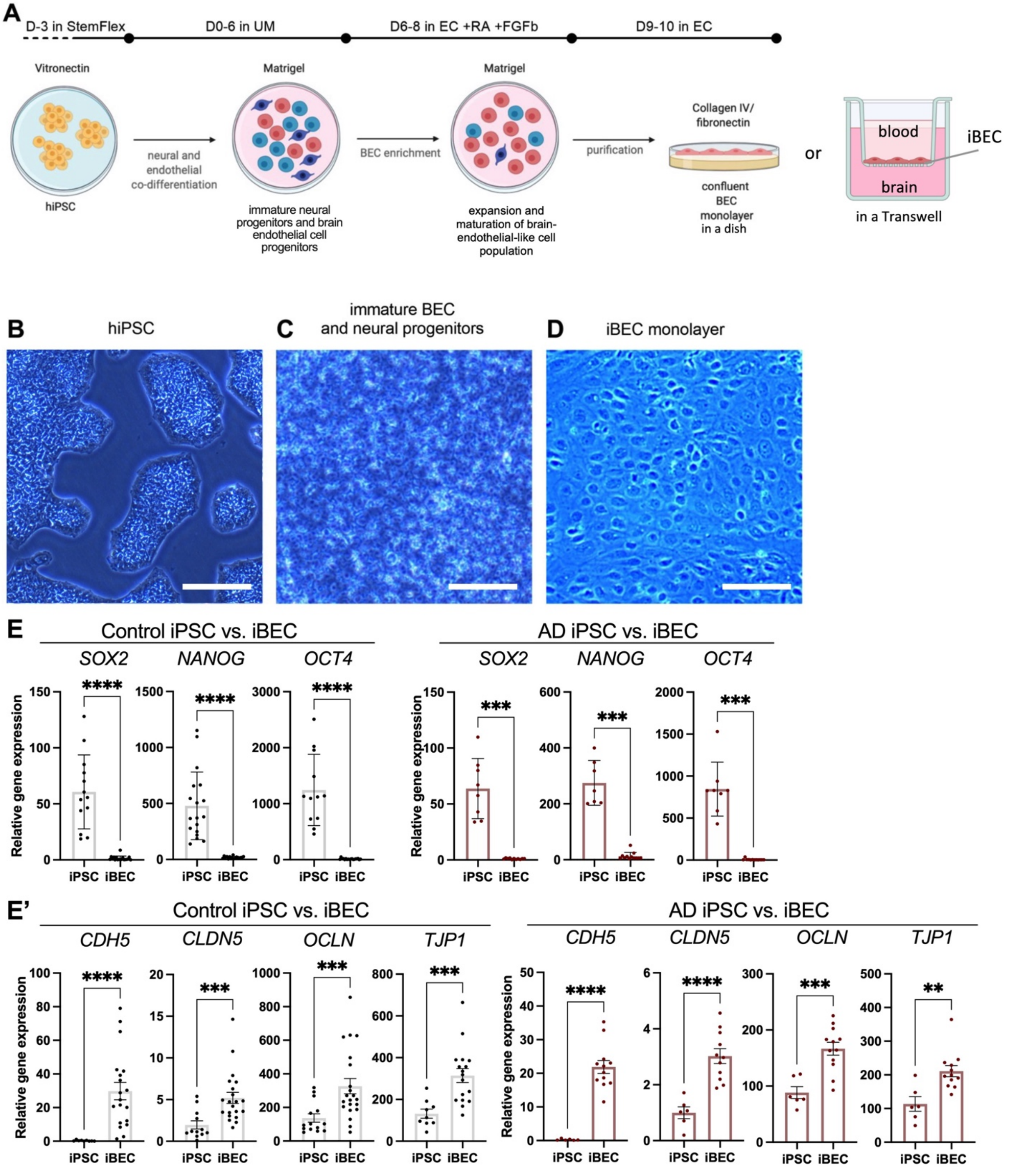

**Figure S3.** (**A**) **Schematic flow of hiPSC-derived iBEC differentiation.** hiPSC are expanded on the vitronectin matrix until 80 % confluency. iBEC differentiation is initiated by plating hiPSC on Matrigel coating and allowed to attach and expand for 3 days. After this, co-differentiation of hiPSC into endothelial and neural progenitors is initiated by switching StemFlex culture media to unconditioned media (lacking FGFb). After 6 days in the unconditioned media (UM), the culture contains a mix of neural progenitors and immature endothelial cells. Immature iBEC are selectively expanded by switching to endothelial cell media (EC), supplemented with retinoic acid (RA) and FGFb. Subsequently, cells are subcultured onto collagen IV/fibronectin matrix, leading to the selective attachment of iBEC and formation of the confluent iBEC monolayer in a cell culture dish or Transwell insert. Phase contrast images of (**B**) hiPSC cultured on vitronectin matrix prior to iBEC differentiation, (**C**) mixed culture containing neural progenitor cells and immature iBEC following 3 days in UM and (**D**) uniform cobble-stone like iBEC monolayer following purification on collagen IV/fibronectin coating. Scale bar, 400 µm in (B), 200 µm in (C) and 100 µm in (D). (**E-E'**) Relative expression of mRNA for (**E**) pluripotency markers *SOX2*, *NANOG* and *OCT4* and (**E'**) brain-endothelial cell marker genes *CDH5*, *CLDN5*, *OCLN* and *TJP1* in control and AD undifferentiated hiPSC and corresponding hiPSC-derived iBEC. Data presented as $\Delta\Delta$CT x $10^6$. Control iBEC: N=3 lines, AD: N=2 lines; a minimum of n=3 independent replicates per line. Data are presented as mean ± SEM. Statistical analysis was performed using unpaired Welch's t-test. *$p$<0.05, **$p$<0.01, ***$p$<0.001, ****$p$<0.0001.

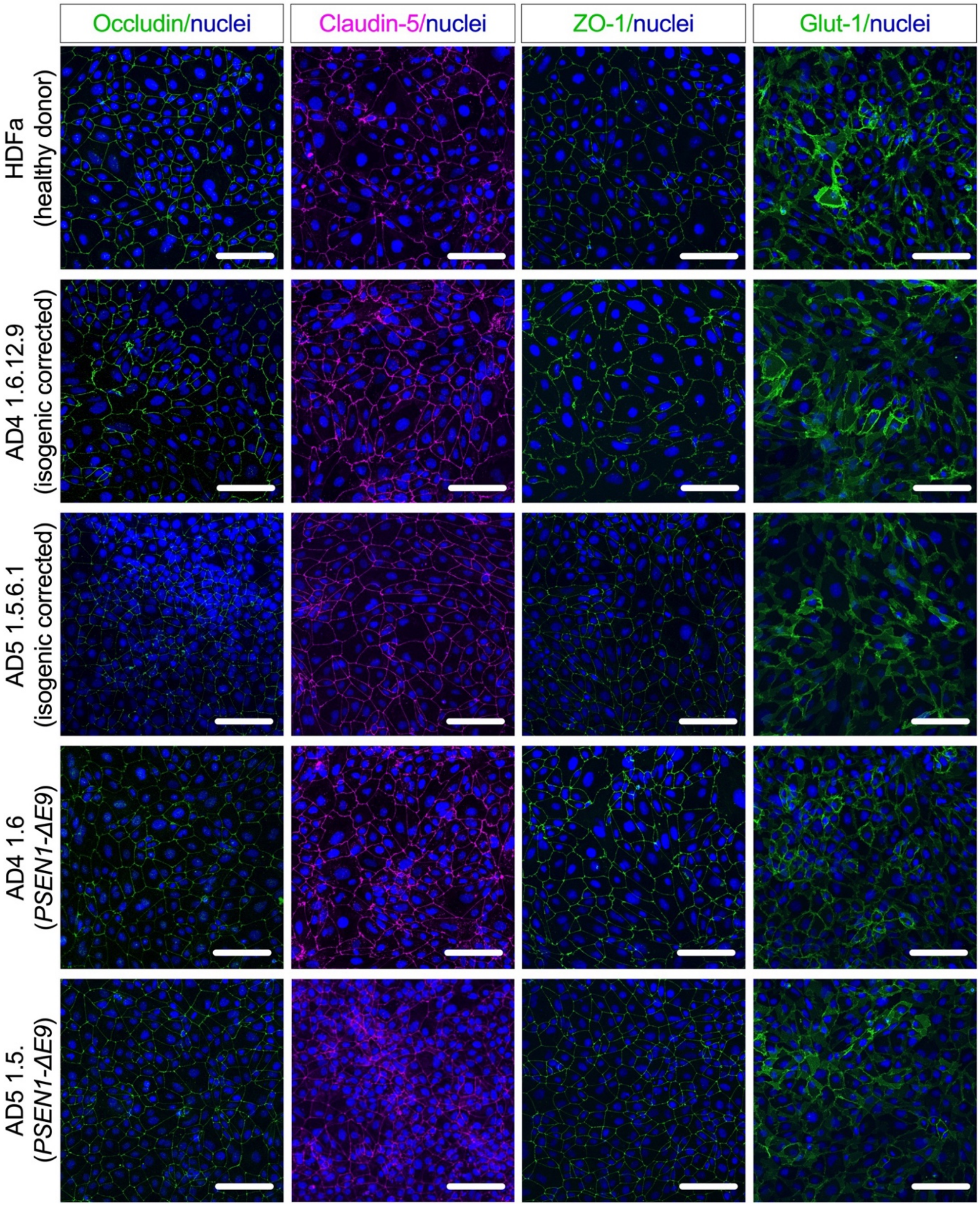

**Figure S4. Characterisation of the healthy donor, isogenic-corrected control and *PSEN1-ΔE9* familial AD iBEC**. Representative immunofluorescence images of brain-endothelial cell markers occludin (green), claudin-5 (magenta), ZO-1 (green) and Glut-1 (green) with Hoechst (blue) nuclear counterstaining in HDFa, isogenic-corrected control AD4 1.6.12.9 and AD5 1.5.6.1, and *PSEN1-ΔE9* AD4 1.6 and AD5 1.5 iBEC. Scale bar, 100 μm.

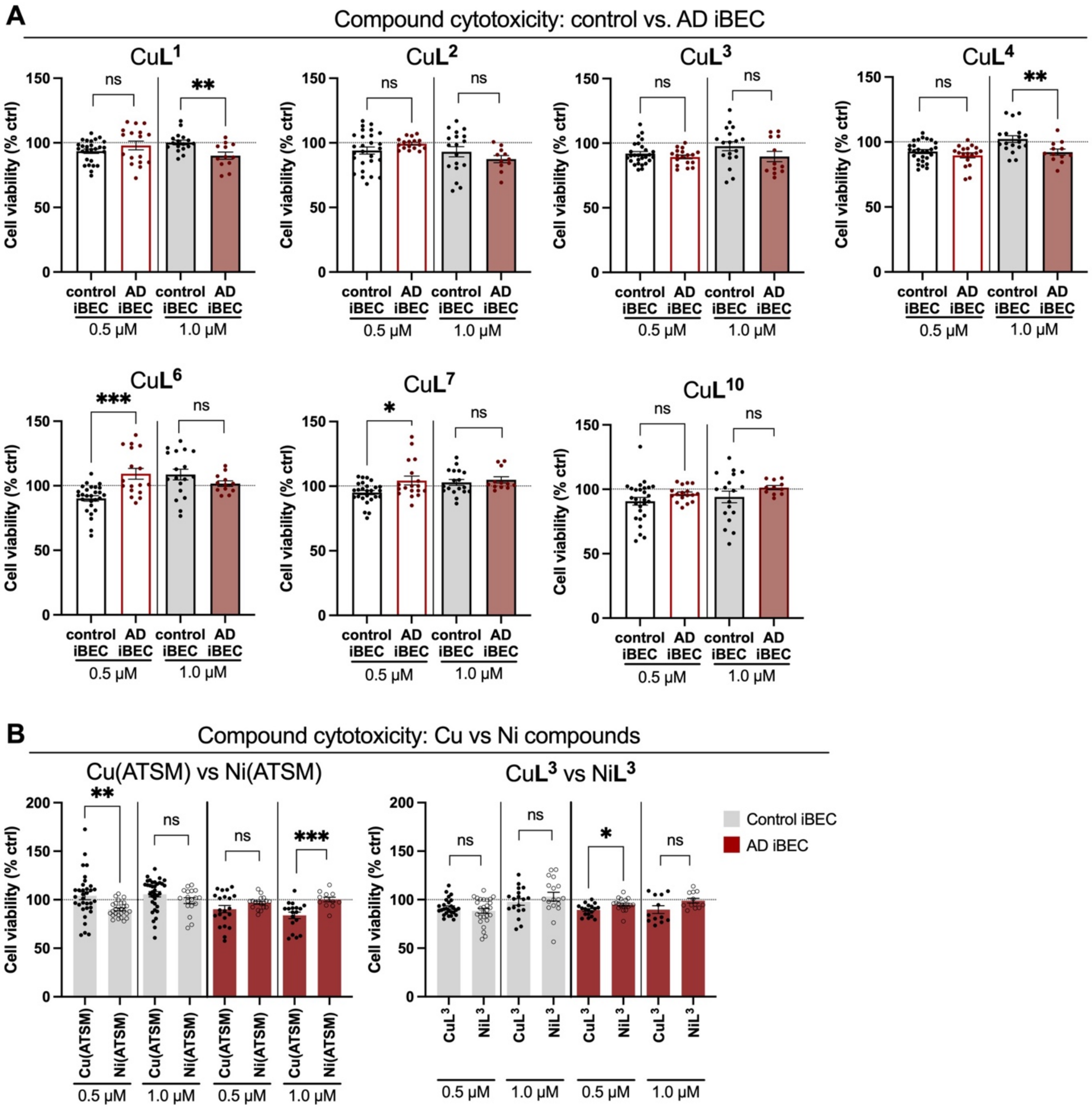

**Figure S5. Comparison of the cytotoxic effects of selected metal compounds between control and AD iBEC.** (**A**) Comparison of control and AD iBEC viability after treatment with 0.5 μM and 1.0 μM of selected Cu and Ni compounds as assessed with MTT assay. (**B**) Comparison of the effect on viability between analogous Cu and Ni analogous compounds in control and AD iBEC. Cell viability is shown as % of viable cells compared to respective untreated control and compared between control and AD iBEC. Control iBEC: N=3 lines, AD: N=2 lines; a minimum of n=5 independent replicates per line per concentration tested. Data are presented as mean ± SEM. Statistical analysis in (A-B) was performed using unpaired Welch's t-test. *$p$<0.05, **$p$<0.01, ***$p$<0.001. The dashed line represents vehicle-treated control.

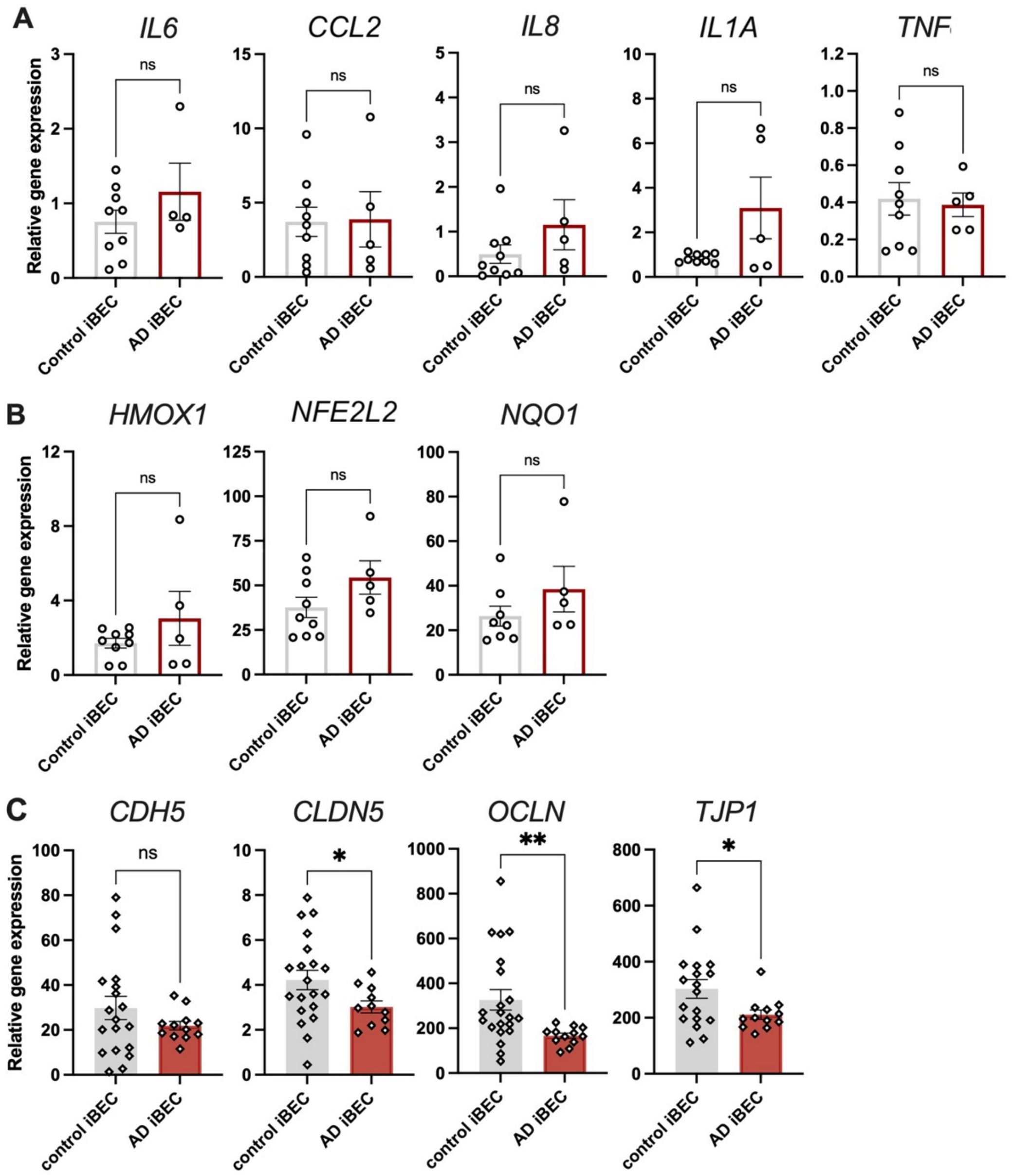

**Figure S6. Baseline expression of proinflammatory, oxidative stress and endothelial cell junctional marker genes in control and AD iBEC**. (**A-B**) Relative expression of mRNA for (**A**) proinflammatory marker genes *IL6*, *CCL2*, *IL8*, *IL1A*, *TNF* and (**B**) oxidative stress-related marker genes *HMOX1*, *NFE2L2* and *NQO1* in control and *PSEN1-ΔE9* AD iBEC. Results presented as $\Delta\Delta$CT x $10^6$. (Control iBEC: N=3 lines, AD iBEC: N=2 lines, minimum n=2 independent replicates per line). (**C**) Relative expression of mRNA for tight and adherens junction marker genes *CDH5*, *CLDN5*, *OCLN* and *TJP1* in control and *PSEN1-ΔE9* AD iBEC. Results presented as $\Delta\Delta$CT x $10^6$. (Control iBEC: N=3 lines, AD iBEC: N=2 lines, minimum n=4 independent replicates per line). Data are presented as mean ± SEM. Statistical analysis in (A-C) was performed using unpaired Welch's t-test. *$p$<0.05, **$p$<0.01.

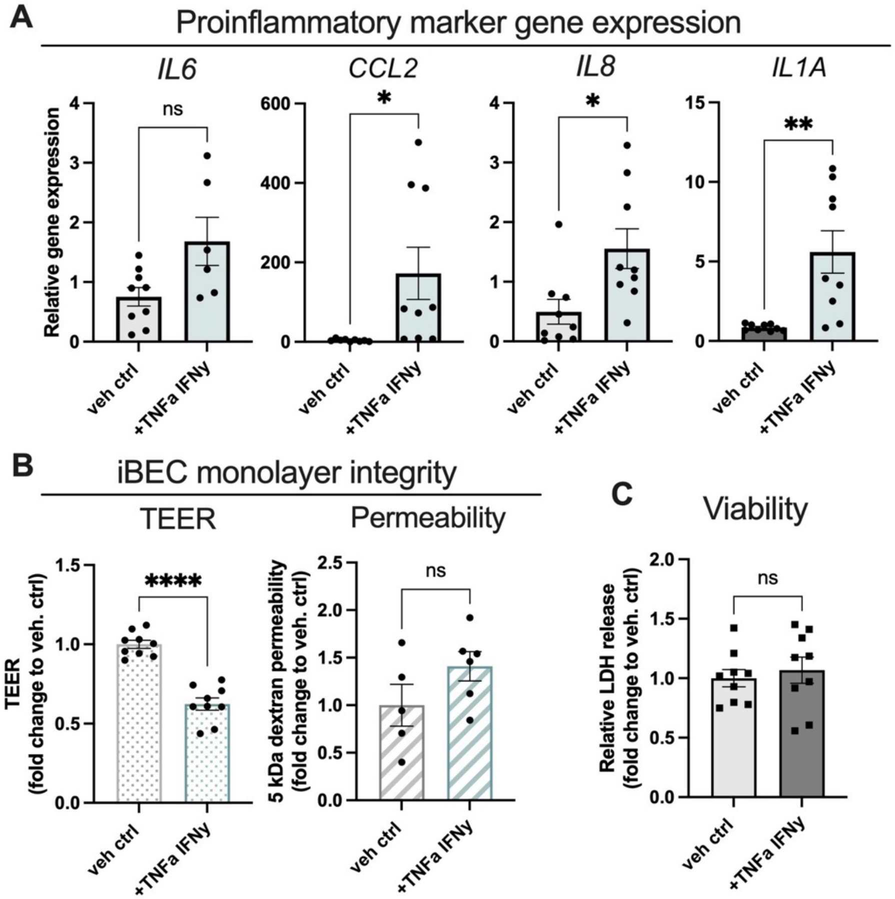

**Figure S7. Effects of TNFα/IFNγ- stimulation on control iBEC phenotype.** (**A**) Relative expression of mRNA for proinflammatory marker genes *IL6*, *CCL2*, *IL8* and *IL1A* in vehicle- and TNFα/IFNγ- treated control iBEC. Results presented as ΔΔCT x $10^6$. (Control iBEC: N=3 lines, n=1-3 independent replicates per line). (**B**) Changes in control iBEC monolayer TEER and passive permeability to 5 kDa dextran following treatment with TNFα/IFNγ. Left panel: data showed as fold change in TEER as compared to vehicle-treated control at 24 h (Control iBEC: N=2 lines, minimum n=3 independent replicates per line). Right panel: data showed as fold change in 5 kDa dextran clearance volume to vehicle-treated control at 24 h (Control iBEC: N=2 lines, n=2-3 independent replicates per line). (**C**) Relative lactate dehydrogenase (LDH) release in control iBEC after stimulation with TNFα/IFNγ. LDH release showed as fold changes to vehicle treated control (veh ctrl). (Control iBEC: N=3 lines, n=3 independent replicates per line). Data are presented as mean ± SEM. Statistical analysis in (A-C) was performed using unpaired Welch's t-test. *$p$<0.05, **$p$<0.01, ****$p$<0.0001. veh ctrl- vehicle-treated control;

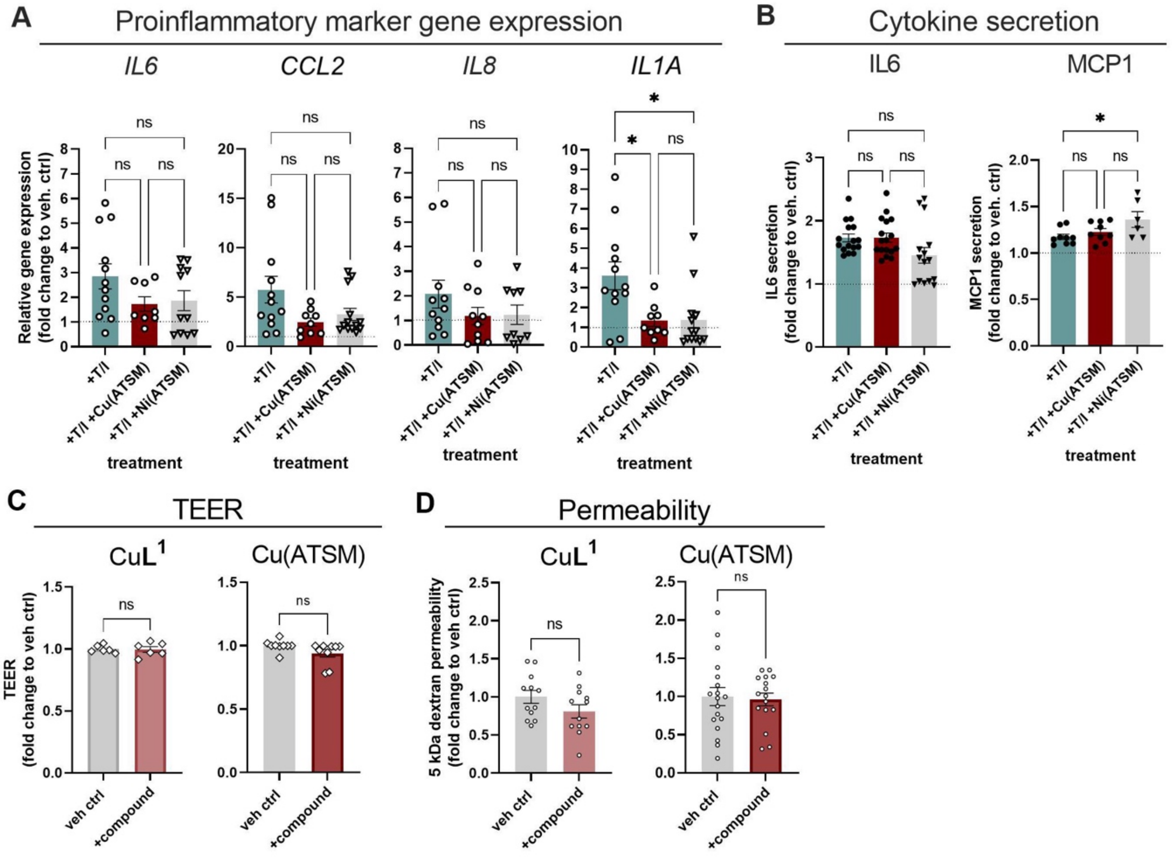

**Figure S8. Comparison of AD iBEC responses to CuL[1], Cu(ATSM) and Ni(ATSM) treatments.** (**A**) Relative expression of mRNA for proinflammatory marker genes *IL6*, *CCL2, IL8,* and *IL1A* in AD iBEC after stimulation with TNFα/IFNγ alone or together with 0.5 μM of Cu(ATSM) or Ni(ATSM) for 24 h. Results presented as fold change in ΔΔCT x $10^6$ as compared to vehicle-treated control (AD iBEC: N=2 lines, minimum n=3 independent replicates per line). (**B**) Secretion of proinflammatory cytokines IL6 and MCP1 in AD iBEC after stimulation with TNFα/IFNγ alone or together with 0.5 μM of Cu(ATSM) or Ni(ATSM) for 24 h. Data showed as [pg/ml] concentration of each cytokine in cell supernatant at 24 h post treatment. (AD iBEC: N=2 lines, minimum n=3 independent replicates per line). (**C**) Changes in AD monolayer TEER following 24 h treatment with 0.5 μM Cu**L**[1] or Cu(ATSM). Data showed as fold change in TEER as compared to vehicle-treated control (veh ctrl) at 24 h. AD iBEC: N=2 lines, minimum n=3 independent replicates per line. (**D**) Changes in AD monolayer permeability to 5 kDa dextran following 24 h treatment with 0.5 μM Cu**L**[1] or Cu(ATSM). Data showed as fold change in 5 kDa dextran clearance volume to vehicle-treated control (veh ctrl) at 24 h. AD iBEC: N=2 lines, minimum n=5 independent replicates per line. Statistical analysis in (A-B) was performed using one-way ANOVA with Tukey's test and in (C-D) using unpaired Welch's t-test. *$p$<0.05. The dashed line represents vehicle-treated control.

**SUPPLEMENTARY TABLES**

Wasielewska *et al*.

**Table S1.** Antibodies used in the study.

| **Primary antibodies** | **Species** | **Source** | **Identifier** |
|---|---|---|---|
| SOX2 | rat | Invitrogen | Cat#14981182 |
| Nanog | rabbit | Abcam | Cat#ab21624 |
| ZO-1 | mouse | Invitrogen | Cat#339100 |
| occludin | rabbit | Invitrogen | Cat#711500 |
| claudin-5 | mouse | Invitrogen | Cat#352500 |
| Glut1 | mouse | Invitrogen | Cat#MA5-11315 |
| **Secondary antibodies** | **Species** | **Source** | **Identifier** |
| anti-mouse Alexa Fluor 488 | goat | Invitrogen | Cat#A11029 |
| anti-mouse Alexa Fluor 594 | goat | Invitrogen | Cat#A11032 |
| anti-mouse Alexa Fluor 647 | goat | Invitrogen | Cat#A32728 |
| anti-rabbit Alexa Fluor 488 | goat | Invitrogen | Cat#A11034 |
| anti-rat Alexa Fluor 647 | goat | Invitrogen | Cat#A21247 |

**Table S2.** Primer sequences used in the study.

| **Target gene** | **Forward primer sequence** | **Reverse primer sequence** |
|---|---|---|
| (Sex determining region Y)-box 2 (*SOX2)* | CCACCTACAGCATGTCC TACTCG | GGGAGGAAGAGGTAA CACAGG |
| Nanog homeobox *(NANOG)* | ACCTCAGCTACAAACAGGT GAA | AAAGGCTGGGGTAGGTAG GT |
| Octamer-binding transcription factor 4 (*OCT4)* | ATCTTCAGGAGATATGCAA AGCAGA | TGATCTGCTGCAGTGTGG GT |
| VE-cadherin *(CDH5)* | AGGCAAGATCAAGTCAAGC GT | GAGTCTCCAGGTTTTCGCC A |
| Claudin-5 *(CLDN5)* | GATTGAGAGGTCTGGGAA GCC | ATCCCATGGCAAACAGAGA GG |
| Occludin *(OCLN)* | GAAGCAAGTGAAGGGATC TGC | ACAACTTGGCATCAGCCTT CT |
| Zonula occludens-1 *(TJP1)* | ACAGCTACAGGAAAATGAC CGA | ACTGGTTCAGGATCAGGA CG |
| Interleukin-6 *(IL6)* | TGCAATAACCACCCCTG ACC | TGCGCAGAATGAGATGA GTTG |

| | | |
|---|---|---|
| Interleukin-8 *(IL8)* | AGACAGCAGAGCACACA AGC | ATGGTTCCTTCCGGTGGT |
| Interleukin-1α *(IL1A)* | CATCGCCAATGACTCAGAG AAG | TGCCAAGCACACCCAGTA GTCTTGCTT |
| Interleukin-1β *(IL1B)* | AATCTGTACCTGTCCTGCGTG TT | TGGGTAATTTTTGGGATCTAC ACTCT |
| Monocyte chemoattractant protein-1 *(CCL2)* | GCTCATAGCAGCCACCTTC ATTC | GGACACTTGCTGCTGGTG ATTC |
| Tumour necrosis factor α (*TNF*) | CAGCCTCTTCTCCTTCC TGAT | GCCAGAGGGCTGATTAG AGA |
| Heme oxygenase 1 (*HMOX1*) | CCCACGCCTACACCCGCT AC | GGTGGCACTGGCAATGTT GG |
| Nuclear factor E2-related factor 2 (*NFE2L2*) | TGCCAACTACTCCCAGGTT G | GACTGGGCTCTCGATGTG AC |
| NAD(P)H quinone dehydrogenase 1 (*NQO1*) | GGGCAAGTCCATCCCAACT G | GCAAGTCAGGGAAGCCTG GA |
| *18S* | TTCGAGGCCCTGTAATTGG A | GCAGCAACTTTAATATACG CTATTGG |

**Table S3**. Coating solution concentration and cell plating density defined per specific culture plate type utilised during iBEC purification step.

| Cell culture vessel | Collagen IV / fibronectin concentration and volume | iBEC plating density |
|---|---|---|
| **96 well plate** | 80 μg/ml and 20 μg/ml<br>100 μl per well | 1 x $10^5$ cell per well |
| **48 well plate** | 80 μg/ml and 20 μg/ml<br>150 μl per well | 3 x $10^5$ cell per well |
| **24 well plate** | 80 μg/ml and 20 μg/ml<br>250 μl per well | 5 x $10^5$ cell per well |
| **Transwell insert, 0.4 μm pore** | 400 μg/ml and 100 μg/ml<br>50 μl per Transwell | 3 x $10^5$ cell per Transwell |
